# Efficient phasing and imputation of low-coverage sequencing data using large reference panels

**DOI:** 10.1101/2020.04.14.040329

**Authors:** S. Rubinacci, D.M. Ribeiro, R. Hofmeister, O. Delaneau

## Abstract

Low-coverage whole genome sequencing followed by imputation has been proposed as a cost-effective genotyping approach for disease and population genetics studies. However, its competitiveness against SNP arrays is undermined as current imputation methods are computationally expensive and unable to leverage large reference panels.

Here, we describe a method, GLIMPSE, for phasing and imputation of low-coverage sequencing datasets from modern reference panels. We demonstrate its remarkable performance across different coverages and human populations. It achieves imputation of a full genome for less than $1, outperforming existing methods by orders of magnitude, with an increased accuracy of more than 20% at rare variants. We also show that 1x coverage enables effective association studies and is better suited than dense SNP arrays to access the impact of rare variations. Overall, this study demonstrates the promising potential of low-coverage imputation and suggests a paradigm shift in the design of future genomic studies.

## Introduction

The design of genome-wide studies in the context of disease and population genetics has drastically changed in the last few years. The reduced cost of next-generation sequencing has allowed the establishment of large-scale high-coverage whole genome sequencing (WGS) projects ^1^ and accelerated the shift from SNP array platforms to next-generation sequencing. However, this shift is not yet fully realised due the still prohibitive cost of high-coverage sequencing for large sample sizes, sometimes in the order of hundreds of thousands samples or even more ^2,3^. In this scenario, low-coverage WGS has been proposed as a cost-effective alternative approach and has been shown to capture the same amount of common variation and more low-frequency variation than standard SNP array platforms ^4^. This is even more pronounced for populations not specifically targeted by commercially available SNP array platforms.

Low-coverage WGS has been already successfully used in the context of Genome-Wide Association Studies (GWAS). For example, the first GWAS using low-coverage sequencing (1.7x) identified two loci associated with major depressive disorder ^5^. Another study showed that 1x WGS was able to find signals missed by standard imputation of SNP arrays ^6^. Recently, a more systematic examination of the power of GWAS based on low-coverage sequencing suggested that 1x sequencing allows discovering twice as many independent significant signals compared to standard SNP arrays imputation ^7^.

Low-coverage sequencing requires a probabilistic representation of the genotypes, in the form of genotype likelihoods, rather than fixed genotype calls. Due to the probabilistic nature of the data and the high missing rates, imputation is used to refine the genotype likelihoods and to fill in the gaps between the sparsely mapped reads by leveraging information from a reference panel of haplotypes. Current methods are not well-suited to achieve this with respect to modern datasets, as they either have a computational time that scales quadratically with the size of the reference panel ^8^, use approximations that result in reduced accuracy ^9^, or are mainly designed to capture common variation for non-human species ^10^.

In this work, we address the challenge of genotype imputation and haplotype phasing of low-coverage sequencing datasets using a reference panel of haplotypes. To this aim, we propose a novel method, GLIMPSE (Genotype Likelihoods Imputation and PhaSing mEthod), that is designed for large-scale studies and reference panels, typically comprising thousands of genomes. We show the remarkable performance of GLIMPSE using low-coverage whole genome sequencing data for both European and African American populations, and we demonstrate that low-coverage sequencing can be confidently used in downstream analyses. We provide GLIMPSE as a part of an open source software suite that makes imputation for low-coverage sequencing data as convenient as for traditional SNP array platforms.

## Results

### GLIMPSE: a new framework for low-coverage sequencing imputation

An overview of the GLIMPSE method is presented in Figure 1. Briefly, the input of GLIMPSE is a matrix of genotype likelihoods computed from low-coverage sequencing data at all variable positions of the reference panel. GLIMPSE refines these likelihoods by iteratively running genotype imputation and haplotype phasing with a Gibbs sampling procedure. At each iteration, a new pair of haplotypes is sampled per target individual by conditioning on haplotypes from the reference panel and from other target individuals.

**Figure 1:**
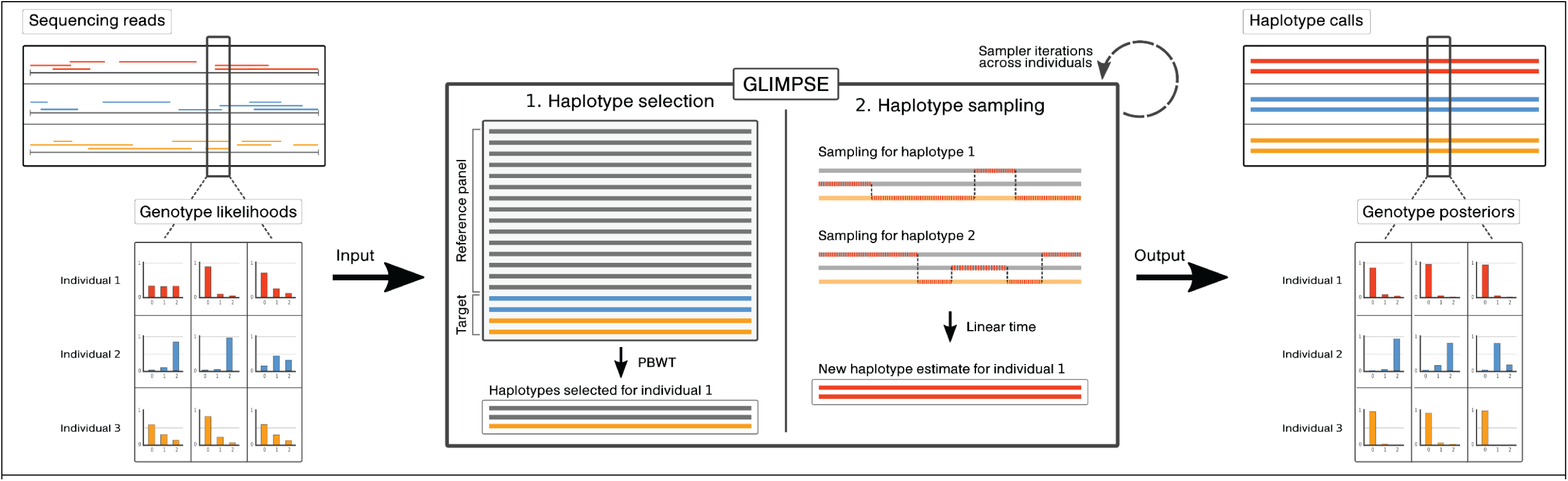
GLIMPSE method overview. The input of the method is a matrix of genotype likelihoods defined at all variable positions obtained directly from the sequencing reads (left). GLIMPSE refines the genotype likelihoods using a Gibbs sampler scheme. At each iteration a new pair of haplotypes for each individual is estimated (middle). This involves two main steps: (1.) the haplotype selection using a reference panel and the current estimate of all other target haplotypes (middle, left) and (2.) a linear time sampling algorithm based on the Li and Stephens model (middle, right). As an output, GLIMPSE produces consensus-based haplotype calls and genotype posteriors at every variable position (right).

To achieve this, GLIMPSE first deconvolves the vector of genotype likelihoods into two independent vectors of haplotype likelihoods, one for each of the two complementary haplotypes (see Online methods). Then, it imputes the two target haplotypes in turn using the resulting likelihoods as additional layers of emission probabilities in a haploid version of the Li and Stephens imputation model ^11,12^. Finally, it updates the phase of the two imputed haplotypes by using the SHAPEIT v4 phasing algorithm ^13^.

The efficiency of this approach stems from two key algorithmic features. First, the model state space is greatly reduced by storing reference and target haplotypes as a Positional Burrows-Wheeler Transform (PBWT) ^14^ which enables the rapid selection of a highly parsimonious and informative subset of conditioning haplotypes to be used in the estimation ^13,15^ (Supplementary Figure 1). Second, the sampling algorithm only relies on computational steps with linear time complexity in the number of conditioning haplotypes being used, which altogether represents a substantial advance over existing algorithms that exhibit quadratic complexity ^8^.

To assess the performance of GLIMPSE in realistic conditions, we used 503 European samples (EUR) and 61 African-American samples (ASW) from the 1,000 Genomes project that have been resequenced at high-coverage (30x) by the New York Genome Center ^16^. From this high-coverage dataset, we called genotypes at all variable positions in the reference panel and used them as validation data in all experiments (Supplementary Table 1). In addition to this, we mimicked typical low-coverage sequencing datasets by downsampling reads to 12 different depths of coverage ranging from 0.1x to 8x and called genotypes for each depth (in the form of genotype likelihoods) at the same set of positions. Finally, we used the Haplotype Reference Consortium (HRC) ^17^ public dataset, which comprises 54,330 haplotypes typed at 39,131,578 variant sites, as the reference panel for imputation.

In all experiments, we assessed imputation accuracy by measuring the squared Pearson correlation between validation and imputed genotypes within minor allele frequency (MAF) bins that we defined from the Genome Aggregation Database (GnomAD v3 ^18^) for the appropriate populations. This accuracy metric is particularly relevant in this context as it quantifies the reduction of effective sample size in downstream association testing due to imperfect imputation.

### Imputation performance of GLIMPSE

We first defined default values for multiple GLIMPSE parameters (Supplementary Figures 2–4) that we used to impute the European samples genome-wide across all simulated sequencing depths (Figure 2A). As expected, imputation accuracy improves as the depth of coverage and the minor allele frequency increase, with most of the differences occurring at rare variants (Minor Allele Frequency; MAF<1%). Remarkably, extremely low-coverages perform well at common variants: 0.3x coverage is highly accurate (r^2^>0.9 for MAF>5%) while 0.1x coverage still provides valuable calls (r^2^>0.7 for MAF>5%). At rare variants, 1x coverage shows accurate results (r^2^=0.8 for MAF=0.1%) while the highest coverage tested, 8x, significantly outperforms all other configurations, as expected (r^2^>0.95 for MAF=0.1%).

**Figure 2:**
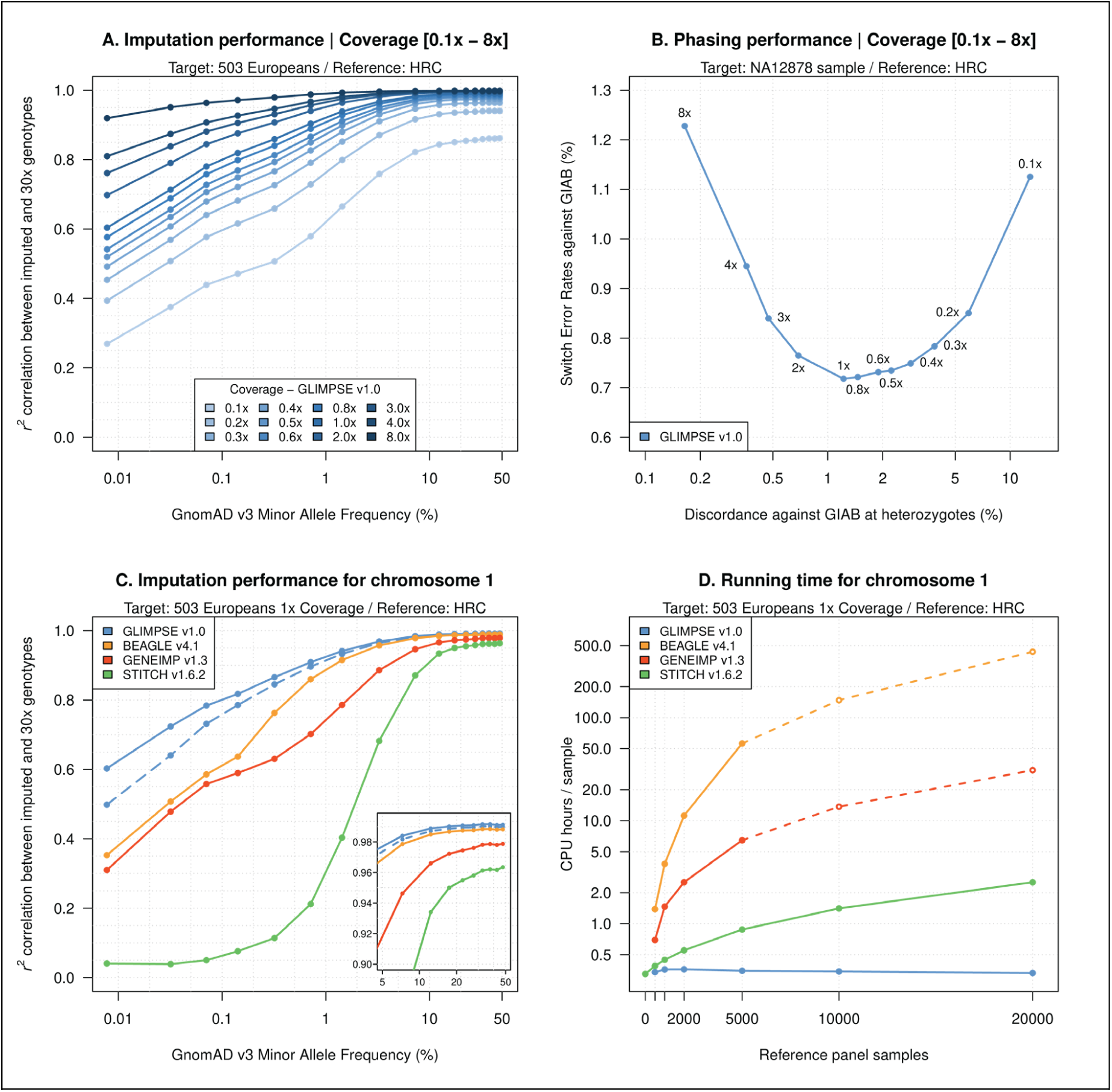
Performance and running time of low-coverage sequencing phasing and imputation. (**A**) Whole genome imputation performance of GLIMPSE at different sequencing coverages of the target dataset. The validation dataset is the 1000 Genomes project samples sequenced at 30x coverage by the New York Genome Center. (**B**) Whole genome phasing performance of GLIMPSE for individual NA12878 while varying the sequencing coverage of the target dataset. We compare the switch error rate (SER, vertical axis) and the discordance rate computed at heterozygous sites (horizontal axis) against Genome In A Bottle (GIAB). (**C**) Imputation performance of low-coverage sequencing methods for the European population chromosome 1 dataset. GLIMPSE was run using 5,000 reference panel samples (dashed blue line) and full reference panel (solid blue line); BEAGLE v4.1 using the 5,000 reference panel samples; GENEIMP v1.3 using 5,000 reference panel samples; STITCH v1.6.2 using the full reference panel. The horizontal axis is on a log scale. (**D**) Running time of low-coverage sequencing methods across different reference panel sizes for the European population chromosome 1 dataset. The vertical axis is on a log scale. Dotted lines represent extrapolated data for the configurations we were not able to run due to time limits.

Overall, we found that even extremely low-coverage sequencing (e.g. 0.3x), followed by imputation from a large reference panel, leads to good assessment of common variants, while at least 1x is required in order to probe rare variants confidently.

Many downstream analyses in population or disease genetics require haplotype level data, which led us to also look at the quality of the haplotypes estimated by GLIMPSE (Figure 2B). For this, we focused on one European sample (NA12878) that has been deeply sequenced and accurately phased using family information and long sequencing reads technologies by the Genome In A Bottle (GIAB) consortium ^19^. When comparing GLIMPSE and GIAB haplotypes, we find that phasing performance varies as a function of the number of heterozygous genotypes and rare variants considered for each tested coverage, thereby forming a convex function. Overall, we obtained low switch error rates ranging between 0.72% and 1.23%, which demonstrates the potential of the haplotypes estimated using GLIMPSE in downstream analyses.

We next compared the performance of GLIMPSE to three well-known low-coverage imputation methods: BEAGLE v4.1 ^8^, GENEIMP v1.3 ^9^ and STITCH v1.6.2 ^10^. To run these methods in reasonable times, we focused on 1x data for chromosome 1, used a randomly downsampled version of the reference panel to 10,000 haplotypes and carefully chose parameter values (Online methods). In contrast, we run GLIMPSE using both the downsampled and full reference panels. The results indicate that our method brings real improvements over other methods across the whole variant frequency range in both situations (Figure 2C, Supplementary Figures 5-10). Indeed, for common variants, GLIMPSE leads to slightly better performance than BEAGLE v4.1 and comfortably outperforms both GENEIMP v1.3 and STITCH v1.6.2. For rare variants the difference is more pronounced, as GLIMPSE greatly outperforms all other methods. For instance, GLIMPSE using the full reference panel provides an accuracy boost of r^2^≈0.2 for variants with a MAF of 0.1%, compared to the second-best method – BEAGLE v4.1. We note that STITCH is primarily designed to impute without the use of a reference panel and would therefore require many more target samples to increase confidence at rare variants.

Notably, the accuracy improvements are obtained in running times that are several orders of magnitude shorter than BEAGLE v4.1 (a hypothetical 1200x speed up when using the full HRC) and match those of STITCH v1.6.2 when run without a reference panel (Figure 2D). A full report of the running times can be found in Supplementary Figures 11 and 12. Moreover, varying the number of reference haplotypes shows that GLIMPSE scales much better than other methods: its running times slightly decrease as the size of the reference panel increases, while BEAGLE scales quadratically and STITCH linearly (Figure 2D; Supplementary Figure 13A). This appreciable property results from the PBWT selection: it finds longer matches in larger reference panels, leading to fewer number of states used in the estimation and therefore smaller running times. Importantly, GLIMPSE running times increase linearly with the number of target samples (Supplementary Figure 13B). Overall, this comparison demonstrates that GLIMPSE brings significant improvements compared to other genotype imputation methods both in terms of accuracy and running times and constitutes the best option available so far to process large amounts of low-coverage sequencing data.

### Imputation performance of low-coverage and SNP arrays

Over the last fifteen years, population and disease genetics studies have routinely been carried out using SNP arrays. In order to show that low-coverage sequencing is a viable alternative to traditional SNP arrays, we assess the performance of genotype imputation performed on low-coverage sequencing and SNP array platforms. For this purpose, we mimicked typical SNP array data by retaining high-coverage genotype calls at sites included on specific SNP arrays. In total, we generated data for 25 different SNP arrays, provided by either Illumina or Thermo Fisher Scientific (Affymetrix) (Supplementary Table 2), that we imputed using two state-of-the-art pipelines (BEAGLE v4.1 ^8^ and v5.1 ^20^) representative of the two last generations of imputation engines. We then compared the accuracy and running times of the various genome-wide imputation strategies, using the HRC as a reference panel. For readability, we only present here a fraction of the results: three different depths of coverage (0.5x, 1x and 4x) and three commonly used SNP arrays (Affymetrix Axiom UK Biobank, Illumina Infinium Omni2.5 and Global Screening Array). The full benchmark is accessible on an interactive web site to facilitate the exploration of the results (Supplementary Figure 14; URL: https://odelaneau.github.io/GLIMPSE/).

We first focused on European samples and found that low-coverage imputation performs well compared to SNP array imputation (Figure 3A). Coverages as low as 1x already outperform dense SNP array models (Illumina Omni2.5) at rare variants (MAF<1%) while matching their accuracy at common variants (MAF>1%). Even at 0.5x coverage, we obtained an appreciable accuracy boost compared to cost-effective SNP arrays such as the Illumina Global Screening Array. As expected, higher coverages, such as 4x, lead to accuracy levels inaccessible to any SNP arrays available on the market, across the full frequency range.

**Figure 3:**
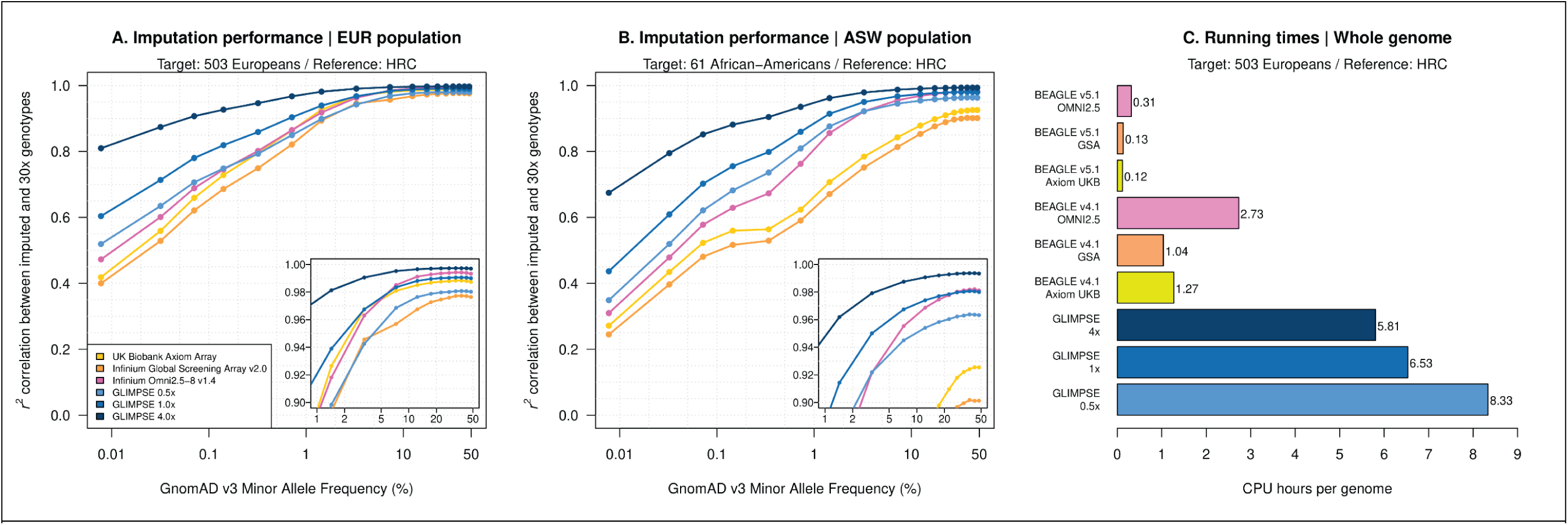
Comparison of low-coverage and SNP array imputation. (**A**) Imputation performance of low-coverage sequencing imputation using GLIMPSE (different coverages) and SNP array imputation using BEAGLE5.1 (different SNP array models) for the European population dataset. (**B**) Imputation performance of low-coverage sequencing imputation using GLIMPSE (different coverages) and SNP array imputation using BEAGLE5.1 (different SNP array models) for the African American population dataset. (**C**) Running time comparison of low-coverage sequencing imputation using GLIMPSE (different coverages) and SNP array imputation using BEAGLE5.1 and BEAGLE 4.1 (different SNP array models).

In the case of African American samples, the improvements brought by low-coverage imputation are even more substantial (Figure 3B). Imputed genotypes at very low-coverage (0.5x) are more accurate than those obtained with standard imputation on the Illumina Global Screening or the Affymetrix Axiom UK Biobank arrays across the full allele frequency range and are only slightly less accurate than imputed genotypes from Illumina Omni2.5 at common variants (MAF > 2%). Higher coverages (e.g. 1x) outperform all three SNP arrays shown for this population. This increased performance in African American samples illustrates the ascertainment bias of SNP arrays towards European populations; a bias absent in low-coverage sequencing.

Overall, we found that 0.5x and 1x sequencing followed by GLIMPSE imputation represents an efficient alternative to sparse or dense SNP arrays, across both European and African American samples. These results are particularly remarkable, given that the SNP array data was simulated under idealistic conditions (i.e. no genotyping errors and low level of missing data). Researchers should therefore consider them as a lower bound on the potential accuracy gain that could be obtained under more realistic conditions.

One caveat of GLIMPSE, compared to standard imputation from SNP arrays, is the higher computational cost involved (Figure 3C). For instance, imputing 1x data using GLIMPSE is 2.4 times slower than imputing Illumina Omni2.5 using BEAGLE v4.1 and 20.8 times slower than imputing the same array using BEAGLE v5.1. However, GLIMPSE imputation remains viable on modern hardware, as imputing a single 1x genome from HRC across ∼40 million variants only requires ∼6.5 CPU hours which corresponds to a cost of $0.65 on an Amazon EC2 m4.large instance ($0.1 per CPU hour).

### Low-coverage sequencing identifies functionally relevant variants

The functional analysis of genetic variation is pivotal in characterizing the genetic architecture of complex traits. In particular, expression quantitative trait loci (eQTL) analyses prove instrumental in this context, by associating genetic variants to gene expression across many individuals. As a proof-of-principle that low-coverage imputation can be confidently used for downstream analyses, we assessed how low-coverage sequencing fares compared to traditional SNP arrays in functional variant analysis.

For this, we mapped eQTLs using the 38 genome-wide call sets generated as part of the previous analysis (1 high-coverage, 12 low-coverages and 25 SNP arrays) together with a RNA-seq dataset of lymphoblastoid cell lines (LCLs) for a subset of 358 European samples ^21^ (Online methods). We first compared the eQTL discovery power across low-coverages and SNP arrays and found that out of 16,894 protein coding and long intergenic non-coding RNA genes tested, between 42.9% and 46.2% associate with genetic variants (eGenes; FDR 5%; Figure 4A). This compares to 46.3% eGenes found when using high-coverage (30x). Notably, sequencing coverages as low as 0.5x can recapitulate most eGenes found with 30x (46.0%) and are matched only by the densest SNP arrays (e.g. Infinium OMNI 2.5; 46.0%). In addition, we confirmed that the set of eGenes and eQTLs detected with most low-coverage datasets closely match those found with high-coverage sequencing. In fact, eGene discovery is matched with 98.1% accuracy for 1x (same as Infinium Omni2.5) and 97.1% for 0.5x coverage (Supplementary Figure 15). In addition, lead eQTLs (the eQTL with the strongest association with the eGene) are perfectly matched in more than 69.3% of the cases for coverages of 1x and above (Supplementary Figure 16A), with association p-values displaying minimal discrepancies (absolute mean errors of the log-transformed p-values of 0.3 for 1x, compared with 0.29 for Infinium Omni2.5; Supplementary Figure 16B).

**Figure 4:**
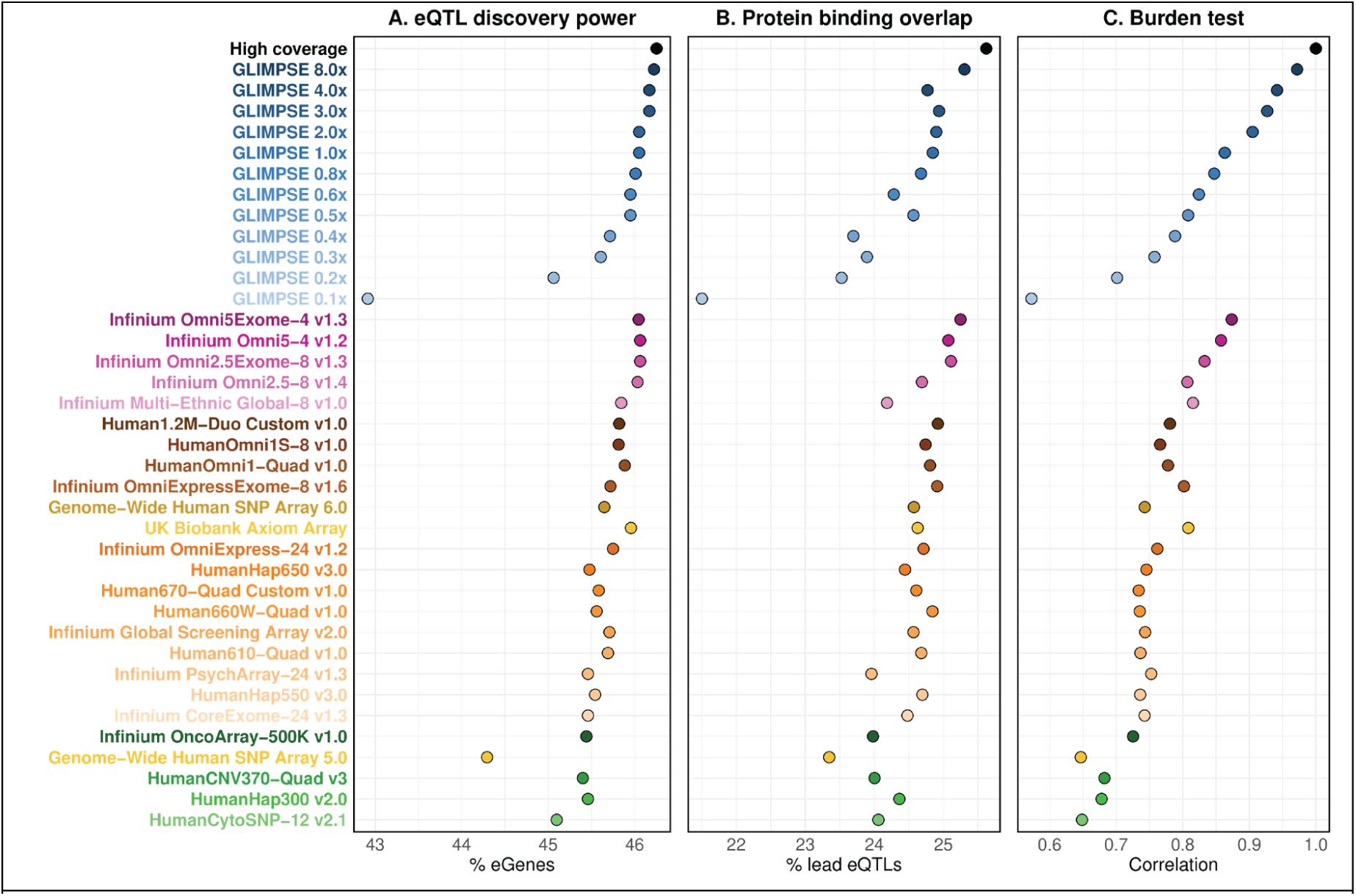
Functional variant analysis across low-coverage and SNP array call sets. (**A**) eQTL discovery power. Percentage of protein coding and lincRNA genes expressed in LCLs whose expression is significantly associated with a variant (eQTL) (FDR 5%). (**B**) Protein binding overlap. Percentage of lead eQTLs overlapping protein binding regions from ChIP-seq experiments performed in LCLs. Only lead eQTLs for eGenes (i.e. significantly associated gene-variant pairs at FDR 5%) were used for each dataset. (**C**) Burden test. Correlation (r^2^) between high-coverage and each assessed low-coverage and SNP array dataset for the number of rare variants (MAF < 1%) found in exons at each protein coding gene. In all panels, SNP arrays are ordered by the number of genotyped variants, from highest (most dense) to lowest.

Previous studies have shown that lead eQTLs often overlap functional genomic regions such as protein binding sites and open chromatin ^22,23^ and this can be used to evaluate the ability of the respective genotyping approaches to pinpoint causal variants in association studies. Here, we found that low-coverage sequencing often identifies lead eQTLs overlapping LCL (i) protein binding regions (derived from ChIP-seq), (ii) open chromatin (DNAse-seq) and (iii) H3K27ac histone modification sites (suggestive of active regulatory elements) at levels comparable to SNP arrays (Figure 4B; Supplementary Figure 17; Online methods). In fact, lead eQTLs identified with 1.0x coverage are found overlapping functional regions as often as those identified with dense SNP arrays such as Infinium Omni2.5. This is a noteworthy result given that only common variants (MAF > 1%) were used in this analysis, as a consequence of the small sample size of the RNA-seq dataset used, and that SNP arrays were attributing perfect genotype calls (no genotype errors introduced) for many of these variants. Importantly, we found that the discovery of functionally relevant lead eQTLs with 0.5x coverage is only slightly lower than high-coverage (e.g. 24.6% for 0.5x, 25.6% for high-coverage in protein binding regions; Figure 4B). As expected, we also confirmed that both discovery power and causal variant mapping improves as coverage increases (Figure 4A-B).

Given that low-coverage sequencing significantly outperforms SNP arrays at rare variants (Figure 3A), we also explored its suitability for burden test analysis of rare coding variants. For this, we used high-coverage sequencing as the ground truth and measured concordance (square Pearson correlation; r^2^) of the minor allele dosages of rare variants (MAF < 1%) overlapping gene exons, at each human protein coding gene (Figure 4C). Notably, we found that low-coverage sequencing performs particularly well in burden test analysis, with coverages above 0.5x having concordances above 0.8 and coverages as low as 1.0x performing as well as the densest SNP array enriched for exonic variants (Infinium Omni 5 Exome). Taken together, these results clearly indicate that sequencing at a low-coverage of 0.5x-1x is well suited to find relevant genetic associations and to assess the potential impact of rare variations, with an accuracy comparable to the most comprehensive SNP arrays.

## Discussion

The accuracy and statistical power boost provided by low-coverage WGS is achieved with the use of genotype imputation from a reference panel of haplotypes, a step aimed at refining uncertain genotype likelihoods and filling in the gaps between sparsely mapped reads. Current methods for low-coverage sequencing imputation are not well-suited for the size of the new generation of reference panels, leading to prohibitive running times and therefore to a drastic reduction in the number of potential applications. In order to alleviate this problem, we present GLIMPSE, a method for haplotype phasing and genotype imputation of low-coverage WGS datasets that reduces the computational cost by several orders of magnitude compared to other methods, allowing accurate imputation based on large reference panels. The efficiency of GLIMPSE is achieved by combining advanced data structures (PBWT ^14^) designed to handle large reference panels with a new powerful linear-time sampling algorithm. As a consequence, the computational time to impute a single variant decreases with the size of the reference panel, an important property since larger reference panels are constantly made available.

We believe that low-coverage WGS provides multiple key advantages over SNP arrays. First, in terms of accuracy as SNP arrays lack a substantial proportion of rare variants and suffer from ascertainment bias; the included variants are biased towards populations involved in the design process of the array ^24^. Low-coverage imputation overcomes both these problems: it provides better typing of low frequency variants independently of the population of interest, without losing power at common variants as shown by our proof-of-principle eQTL analysis. Second, in terms of study design, as the choice of the SNP array requires considering the density of the array and the population of interest. In contrast, only the sequencing depth needs to be considered in the case of low-coverage WGS. Finally, in terms of financial cost; it has recently been showed that sequencing a genome at 1x coverage is at least as expensive as using a sparse SNP array (e.g. Illumina HumanCoreExome array) or half the price of a dense SNP array (e.g. Illumina Omni2.5) ^7^. So far, this financial advantage has been undermined by the prohibitive computational costs of the previous generation of imputation methods. GLIMPSE solves this problem by performing imputation in a cost-efficient manner: a genome sequenced at 1x coverage is imputed from the HRC reference panel for less than $1 on a modern computing cluster. Even though this is one order of magnitude higher than standard pre-phasing and genotype imputation pipelines for SNP arrays, this remains a small fraction of the total cost involved in low-coverage WGS. Overall, our results suggest that whole-genome sequencing at 1x coverage followed by GLIMPSE imputation constitutes an efficient alternative to the densest SNP arrays in terms of cost and accuracy.

We have shown that GLIMPSE already represents an efficient solution for the imputation of low-coverage WGS. However, we think that further method developments are still possible. Indeed, GLIMPSE relies on the existence of a reference panel and on genotype likelihoods computed from the sequencing reads. While reference panels are already available for multiple human populations, this is not always the case and we think that our model can easily be extended to perform imputation without a reference panel. Additional efforts are also required to improve genotype likelihood calculation and data management. This involves the development of specific file formats to store genotype likelihoods, which could exploit the sparsity inherent to low-coverage datasets for data compression. Furthermore, running times can be further decreased by implementing techniques successfully adopted in the context of SNP array imputation, such as delayed imputation ^20^ and the use of compressed file formats for reference panels ^15,20^. This might further reduce the computational gap between the two types of imputation approaches.

As next-generation sequencing is getting more economically sustainable, the use of low-coverage sequencing for future genomic studies will probably become more popular in the near future. This has not been possible so far, mainly due to the lack of efficient computational methods able to efficiently impute this type of data. In order to facilitate this transition, we provide a complete open-source software suite that covers the full process of phasing and imputation of low-coverage sequencing datasets for a wide range of applications in genetics.

## Online Methods

### Datasets

#### NYGC 1000 Genomes Project data

The 1000 Genomes Project ^16^ is a landmark project in human genetics. It was designed to provide a comprehensive description of genetic variation across different ancestries and it has been used in a wide range of studies in human genetics. The phase 3 panel contains sequenced data of 2,504 individuals sampled from 26 different populations that can be divided into five continental groups: 661 samples with African ancestry (AFR), 347 samples with American ancestry (AMR), 503 samples with European ancestry (EUR), 504 samples with East Asian ancestry (EAS) and 489 samples with South Asian ancestry (SAS). Recently, the New York Genome Center (NYGC), funded by National Human Genome Research Institute, has resequenced all 2,504 samples at 30x depth of coverage. This dataset is available on the European Nucleotide Archive (ENA; accession PRJEB31736). Sequencing was performed using an Illumina NovaSeq 6000 system with 2×150bp reads that were aligned to GRCh38. Detailed description of the pipeline used to process the sequencing data can be found on the EBI FTP server (<u>link</u>). In the context of this work, we only used the sequence data available for 503 European (CEU, GBR, TSI, IBS and FIN) and 61 African-American (ASW) individuals.

#### Haplotype Reference Consortium

The Haplotype Reference Consortium (HRC) ^17^ combines sequence data across 32,470 individuals from multiple cohorts with low-coverage WGS (from 4x to 8x coverage) of subjects with mostly European ancestry. The dataset contains a set of 39,235,157 variants (bi-allelic SNVs) with minor allele counts (MAC) ≥ 5, collected from 20 different studies. The publicly available version of the data contains 27,165 individuals (chromosomes 2-22, X) and 22,691 individuals (chromosome 1), with phased SNV genotypes coded in the Human genome assembly GRCh37. This version of the HRC data was downloaded from the European Genome-Phenome Archive at the European Bioinformatics Institute (accession EGAS00001001710). The picard toolkit (URL: http://broadinstitute.github.io/picard/) was used to perform the liftover of the data to the Human genome assembly GRCh38. We discarded data for strand flips, obtaining on average across all the chromosomes that 99.8% of the variants were conserved after the liftover.

We used the HRC reference panel in all the imputation tasks we performed, for both the European and African American imputation experiments. In each of these cases, we removed the target population from the reference panel, since the HRC dataset contains 1000 Genomes samples. We used the full reference panel for all the results shown in Figures 3-4 and Supplementary Figures 2-4,13. For the results shown in Figure 2 and Supplementary Figures 5-12, we randomly downsampled the HRC data on chromosome 1 in order to form smaller reference panels containing 20,000, 10,000, 5,000, 1,000 and 500 samples ensuring that HRC_500_ ⊆ HRC_1000_ ⊆ HRC_2000_ ⊆ … ⊆ HRC_20000_. We kept monomorphic sites in each of the downsampled datasets to maintain the exact same number of variants.

#### Genotype likelihoods computation

Genotype calling from the NYGC sequencing data, either downsampled or not, was performed using the following procedure. First, we extracted all variable positions in the HRC together with the reported reference and non-reference alleles. Then, we run bcftools mpileup and call modes on the CRAM files specifying the exact positions and alleles at which to perform genotype calling (see GLIMPSE online documentation for the exact set of command lines). Briefly, (i) all PCR duplicates as annotated by Picard have been excluded from the analysis, (ii) the coverage was capped to a maximum of 250 per position and (iii) reads having a mapping quality above 0 and a base quality above 13 were kept for the calling.

As a result of this, we enforced the computation of genotype likelihoods at all the 39,235,157 bi-allelic SNVs present in HRC. For all the low-coverage imputation experiments, we used all variant sites and genotypes, regardless of the certainty of the likelihoods. For the validation data based on high-coverage calls, we only used the genotypes that were derived from at least 8x coverage and exhibiting high certainty of the call given the genotype likelihoods. To do so, we assume uniform prior for all three possible genotypes (Ref/Ref, Alt/Ref and Alt/Alt), computed the genotype posteriors from the likelihoods and the prior, and only kept in the validation the genotypes having a posterior probability above 0.9999. Since genotype calls have also been made by the NYGC using a different pipeline based on GATK, we checked the concordance between our high-coverage calls and those made by the NYGC and found a discordance rate below 0.01% (1 discordance out of 10,000 genotypes). This level of discordance matches well what is expected given the threshold we used on the posteriors (i.e. 0.9999) and demonstrates the high quality of the genotype calls produced by our pipeline. All genotype likelihoods we produced were phred scaled and stored in VCF/BCF files using the FORMAT/PL field.

#### Simulating SNP array data

We simulated SNP array data of a wide range of 25 different SNP array models produced by either Illumina or Thermo Fisher Scientific (Affymetrix), from high-quality genotype calls obtained from NYGC 1000 Genomes 30x data. To do so, we first downloaded the lists of variable positions included in the SNP arrays of interest from https://www.well.ox.ac.uk/~wrayner/strand/ mapped on the reference human genome GRCh38. We used the validation data described above to obtain genotype calls at these specific positions for all target individuals (i.e. posteriors > 0.9999 and coverage >= 8x). Variant sites not included in HRC or having a missingness greater than 5% were removed from the respective SNP array datasets (Supplementary Table 2). We developed a program in the GLIMPSE tools suite (GLIMPSE *snparray*) to achieve this specific task.

We remark that the simulated SNP array data has been generated under an idealistic scenario since (i) no genotyping errors were introduced (i.e. genotypes perfectly match those in the validation data) and (ii) the quality of the genotypes does not depend on minor allele frequency, which is typically the case for standard SNP arrays: low frequency variants are more challenging to accurately call ^25^.

#### Geuvadis RNA-seq data

We used BAM files previously mapped to GRCh37 for RNA-seq experiments on lymphoblastoid cell lines (LCL) from a subset of 358 European (EUR) individuals from the Geuvadis study ^21^ also present in the 1000 Genomes project phase 3. Data was downloaded from the EBI ArrayExpress (accession code E-GEUV-1). We quantified gene expression for protein-coding and long intergenic non-coding RNAs (lincRNAs) as annotated in GENCODE v19 annotations ^26^ using QTLtools v1.1 ^27^. We excluded from the analysis all genes within or around the MHC complex region (chr6:29500000-33600000) and in non-autosomal chromosomes (X and Y), as well as those with poor variability across individuals (>=50% of the individuals with RPKM=0). We liftovered gene coordinates from GRCh37 to GRCh38 using the UCSC liftover tool, determining a final set of 16,894 genes left for the eQTL analysis. To account for confounding factors, we regressed out the following covariates: sex, ancestry (3 first PCA principal components (PC) computed on the 30x coverage genotype data) and technical variables (50 PCA PCs detected on the phenotype data, determined as the number of PCs that maximizes the number of eGenes discovered). We finally normalized the expression quantifications across individuals to match a normal distribution *N*(0,1).

### GLIMPSE model for low-coverage sequence data

#### Notation and genotype likelihoods

GLIMPSE is a method for haplotype phasing and genotype imputation for a set *T* containing *Q* unrelated individuals. The input of the algorithm is primarily made of two components: a reference panel of haplotypes *H* and a matrix of genotype likelihoods (GLs). A typical reference panel for GLIMPSE contains data for at least thousands of samples and millions of variants, as those provided by consortia such as the 1000 Genomes ^16^ or the Haplotype Reference Consortium ^17^. The matrix of GLs is computed using standard variant calling pipelines from the available low-coverage sequencing data. A genotype likelihood is defined as the probability of observing the sequencing reads at a specific genomic position given the unknown underlying genotype. This probability distribution is therefore uniformly distributed when no reads are available and gets peaked towards a specific genotype as the number of observed reads supporting the genotype increases. In the context of this work, genotype likelihoods are computed at all variable positions reported in the reference panel, which implies that no variant discovery step is needed. In practice, we achieve this using bcftools ^28^, but Genome Analysis Toolkit (GATK) ^29^ can also be considered for this task.

Let us denote the reference panel as *H*, typed at *M* bi-allelic variants. We consider a single sample *i* in the set of target samples *T*. From the sequencing reads of the individual *i*, we can compute the genotype likelihood in all the *M* bi-allelic variants, indicating the probability of observing the sequencing reads *R*_*i*_ given the unobserved genotype *g*_*i,j*_. The genotype likelihood for an individual *i* at site *j* is a three-dimensional vector *GL*_*i,j*_ = *Pr*(*R*_*i*_|*g*_*i,j*_), where *g*_*i,j*_ ∈ {*rr, ra, aa*} indicates the reference homozygous, the heterozygous and the alternative homozygous genotypes, respectively. There are a range of methods for genotype likelihood calculations, that use the mapping and quality scores of reads at a given site. We consider the case where genotype likelihoods are computed for the full set of target samples *T* each variant of the reference panel *H*, assuming uniform likelihood distributions in case of missing information.

GLIMPSE makes use of genotype likelihoods to run haploid imputation. For this reason, GLIMPSE derives a haplotype likelihood distribution from the genotype likelihood distribution by conditioning the genotype likelihoods on the current estimate of one of the two haplotypes for the sample.

Let us indicate a pair of haplotypes for the sample *i* as 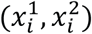, and 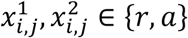 be reference and alternative alleles at the site *j*, respectively. Then, by fixing the haplotype 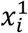 we can derive the haplotype likelihood for haplotype 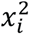 at site *j* as a two-dimensional vector 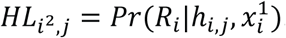. Conversely, 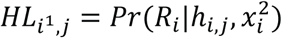 indicates the haplotype likelihood of the first haplotype at locus *j*, fixing the value of the second haplotype.

At the start of the GLIMPSE algorithm, no phase information is available. In this case, the haplotype likelihoods are initialized as:

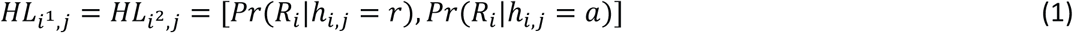

where:

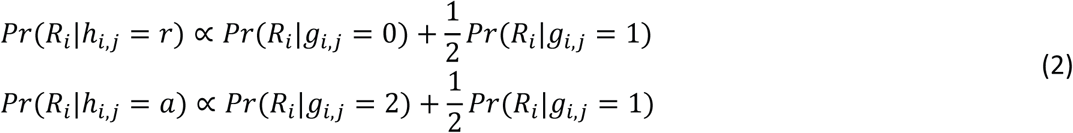

When the haplotype likelihood is conditional on the phase of the other haplotype (e.g. 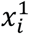), this becomes:

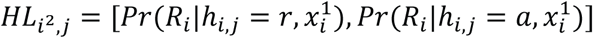

By fixing the value of one haplotype, one of the possible genotype configurations becomes invalid and this reduces to:

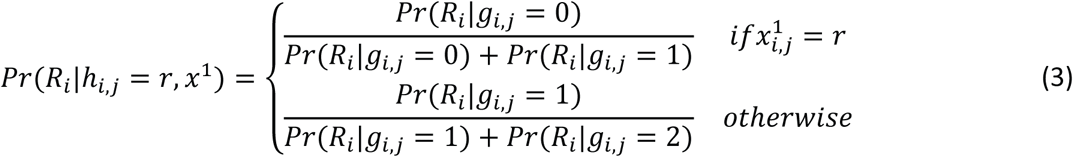

Similarly, we can compute 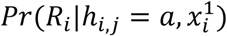. For convenience, we use the notation 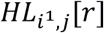 and 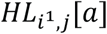 to indicate the reference and alternative haplotype likelihood for the first haplotype of the *i*th individual at the *j*th locus.

#### Model description

GLIMPSE implements a Gibbs sampling procedure, extending the SHAPEIT v4 model. At each iteration, the GLIMPSE estimates a consistent pair of haplotypes 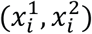 for each sample *i*, based on its genotype likelihoods, the reference panel of haplotypes *H* having 2*P* haplotypes and the 2*Q* − 2 haplotypes previously estimated for the other target individuals.

Let us only consider a single iteration *n*, where we aim to sample a new pair of haplotypes 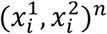 for the individual *i*, and let us indicate the previous haplotype pair for the same individual as 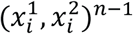.

Let us assume that the reference panel plus the current haplotypes estimate have a total of *K* = 2*P* + 2*Q* haplotypes. The state selection algorithm based on this set returns a subset of size 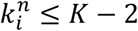 haplotypes on which haploid imputation is performed. Let us denote this set as 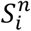 and the set of states retrieved in the previous iteration as 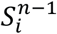 having size 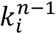.

The first step of a new iteration is to refine the phasing of the diplotype generated in the previous iteration, 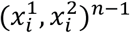 by sampling a new estimate. For this task we use the SHAPEIT v4 model, which divides the region in several consecutive non-overlapping segments, such that each segment contains eight possible haplotypes consistent with the current genotype (three heterozygous positions). Given the segment representation, sampling a pair of haplotypes given a set of known haplotypes 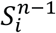 involves sampling from the posterior distribution 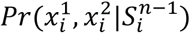 and exploiting the segmentation this procedure has linear time complexity with the conditioning states 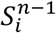. Detailed description of the sampling model has already been published and can be found in ^13^.

After a new diplotype has been generated, the next few steps aim at improving the quality of the genotypes by performing a step of haploid imputation on both the newly generated haplotypes. Before the haploid imputation step, we generate a new set of conditioning states 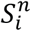 on which haploid imputation will be performed.

The imputation procedure involves: (i) update the haplotype likelihoods of the target haplotype based on the most updated value of the other haplotype in the pair, (ii) run a modified version of the haploid Li and Stephens HMM where the emission layer takes into account the haploid likelihoods and (iii) update the value of the haplotype by sampling from the posterior distribution generated in step (ii).

Summarising, at *n*th iteration the algorithm performs the following steps for the individual *i*:

1. Runs a diploid phasing step, starting from 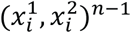 in order to get a new estimate 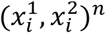;
2. Reduces the state space to new subset 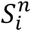, using the new estimate 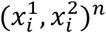;
3. Computes haplotype likelihoods for 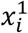 conditional to 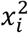, getting 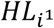;
4. Runs a haploid phasing step for the haplotype 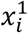, getting haploid posterior probabilities.
5. Updates 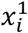 by sampling from the haploid posterior distribution;
6. Computes haplotype likelihoods for 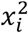 conditional to 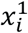, getting 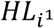;
7. Runs a haploid phasing step for the haplotype 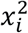, getting haploid posterior probabilities.
8. Updates 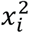 by sampling from the haploid posterior distribution;

#### Haploid imputation

The haploid imputation step of the model relies on a modified version of Li and Stephens hidden Markov model used by several imputation methods ^8,12^. Let us assume we want to perform haploid imputation for the first haplotype *i*-th sample, called 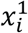. We also assume we have computed the haplotype likelihoods 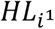 for haplotype 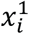, based on the value of 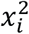 as well as the set of conditioning states *S*_*i*_ of size *k*.

The probability of observing 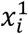 from *S*_*i*_ can be then written as:

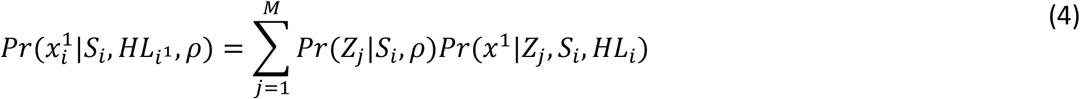

where *Z* = {*Z*_1_, *Z*_2_, …, *Z*_*M*_} is a sequence of unobserved copying labels defined in the space of *S*_*i*_, *Z*_*j*_ ∈ *S*_*i*_. Since *Z*_*j*_is a label over the states *S*_*i*_, we indicate as *S*_*i*_[*Z*_*j*_, *j*] the haplotype value for the state indicated by *Z*_*j*_ at marker *j*. The vector *ρ* = {*ρ*_1_, *ρ*_2_, …, *ρ*_*M*−1_} is a parameter modelling recombination event.

The two terms of equation (4) model transition and emission probabilities of the HMM. The transition probability *Pr*(*Z*_*j*_|*S*_*i*_, *ρ*) is defined as:

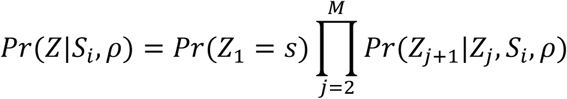

With:

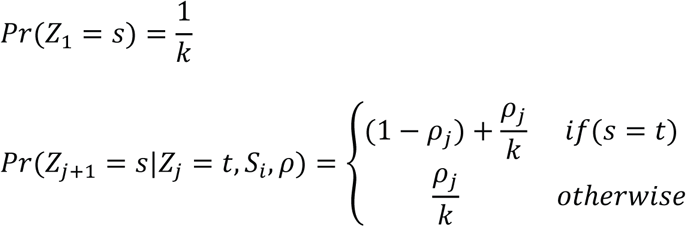

where 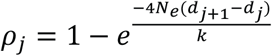, where *N*_*e*_ is the effective diploid population size, (*d*_*j*+1_ − *d*_*j*_) is the genetic distance between marker *j* + 1and *j*, and *s, t* are two possible values of *Z* defined as categorical label in the space *S*_*i*_.

The emission probability in GLIMPSE is modified compared to the standard Li and Stephens HMM. We use *μ* = 10^−4^ as the probability of genotyping error at each considered site. The emission probability can be described as follows:

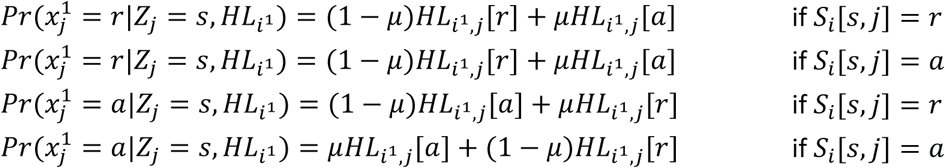

#### Initialization and Gibbs iterations

The GLIMPSE iterations scheme is formed by three main parts: (i) an initializing iteration (ii) a set of burn-in iterations to guarantee convergence and (iii) a set of main iterations.

GLIMPSE starts with an initialising iteration to provide a first haplotype estimate. It does so by selecting K (K=1,000 by default) random haplotypes in the reference panel, but it is also possible to provide a list of samples in the reference panel from which initializing haplotypes are chosen. In this way it is possible to select haplotypes genetically similar to the target population. When the set of haplotypes has been selected, two haploid imputation steps are performed, one for each haplotype. Haplotypes are then sampled using posteriors given by imputation.

In the burn-in and main iterations the first step is diploid phasing, given the previous haplotype estimates. This step is performed prior to the state selection and this is crucial to get accurate states. The diploid phasing scheme is a simplified version of the SHAPEIT v4 approach ^13^. After the phasing iterations, the two haploid imputation steps are performed, one of each haplotype. The only difference between burn-in and main iterations regards the fact that output is only stored in the main iterations. GLIMPSE averages genotype imputation posteriors across multiple iterations. This feature makes GLIMPSE resistant to noise and parameter settings as shown by (Supplementary Figure 2-5). Phasing information of the haplotype pairs sampled throughout the main iterations are recorded in the output HS field in the VCF file and can be used by the GLIMPSE sampling algorithm to provide accurate consensus haplotype calls.

#### State selection

GLIMPSE reduces the state selection space by creating the PBWT of the reference panel and the current estimate of the target haplotypes, in order to identify a subset of haplotypes that share long identity by state (IBS) sequences with each of the target haplotypes. The state selection is performed before the diploid sampling and haploid imputation steps.

As performed in SHAPEIT v4 ^13^ and IMPUTE v5 ^15^ models, the PBWT is constructed in the standard order from left to right across the region and the state selection occurs at specific markers (defined by the pbwt-modulo parameter), during the construction.

GLIMPSE implements a PBWT selection scheme adopted by IMPUTE v5, called neighbour selection algorithm. The PBWT of the reference panel and the current estimated haplotypes is created. By exploiting the fact that longest reverse prefixes are by definition in the neighbourhood of the target haplotype position in the PBWT, at every selection marker (defined by the pbwt-modulo parameter), the algorithm simply select the set of 2L neighbouring states (L in both the directions) from the current haplotype position and stores them in a list. This procedure is repeated for each target haplotype. When the copying list of all the target haplotypes has been updated, the PBWT of the next marker is computed. Finally, at the end of the PBWT step, a single set of selected states is created for each target sample, by merging the lists of copying states retrieved by both the sample’s haplotypes. A schematic illustration of the selection algorithm is shown in (Supplementary Figure 1).

The number of states can be controlled using the pbwt-modulo (default pbwt-modulo=8) and pbwt-depth (pbwt-depth=2) options, referring to the frequency of the PBWT selection (in number of variants) and the number of neighbouring states used for the selection at a specific marker, respectively. Incrementing the pbwt-depth parameter and decreasing the pbwt-modulo parameter has the effect of increasing the number of states retrieved, therefore gaining accuracy at the cost of more computational time.

The number of states cannot increase arbitrarily in GLIMPSE, because it is capped by the number of states used for initialization (K=1,000 by default). In the case the number of states retrieved exceeds the limit, states are sampled from the frequency distribution of appearance in the selection.

A striking feature of the state selection algorithm is that the number of selected states decreases as the number of haplotypes of the conditioning set increases. This property allows GLIMPSE to run the costliest part of the algorithm (diploid phasing and haploid imputation) on a small subset of states that decreases when the reference panel increases. For this reason, the cost per sample shown in (Figure 2D, Supplementary Figures 11-12) slightly decreases when increasing the number of haplotypes in the reference panel.

#### File formats

GLIMPSE requires both the input genotype likelihoods and reference haplotypes to be in indexed VCF/BCF format. For the reference panel the FORMAT/GT field is used, and phased haplotypes are required. For the target panel only phred-scaled likelihoods are read in the FORMAT/PL field. GLIMPSE reads only biallelic variants defined in both datasets.

All variants present in the reference panel are assumed to be present in the target dataset containing genotype likelihoods. In the case of completely missing data, a flat likelihood is called by GLIMPSE, allowing the method to perform standard imputation. This shows how GLIMPSE generalises standard imputation, dealing with completely missing information as part of the model, instead of requiring additional features. Indeed, standard SNP array can be viewed as a special case of low-coverage sequencing imputation, where the likelihoods have either no uncertainty (positions called in the SNP array) or full uncertainty (missing data, resulting in a flat distribution of the genotype likelihood).

#### Phred-scaled genotype likelihoods

GLIMPSE requires as an input the normalized phred-scaled genotype likelihood for each variant and individual to be imputed. The phred-scaled likelihoods are recorded in the FORMAT/PL columns of the target VCF/BCF file. The phred-scaled genotype likelihoods are a logarithmic transformation of the three-dimensional vector *GL*_*i,j*_ defined as *PL*_*i,j*_[*g*_*i,l*_] = −10log(*Pr*(*R*_*i*_|*g*_*i,j*_)), where *g*_*i,j*_ ∈ {*rr, ra, aa*} indicates the reference homozygous, heterozygous and alternative homozygous genotype, respectively. A normalisation is then applied, so that the lowest PL value of the three-dimensional vector is set zero and the others normalised accordingly.

From the normalised likelihoods, it is possible to get the original normalized posterior probabilities by simply converting them to real space, then normalising them. In practice:

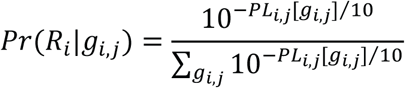

For efficiency reasons, GLIMPSE unphread the phred-scaled genotype likelihoods by using a pre-computed discretized map that allows a fast computation of the exponential 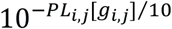.

#### Imputation chunks and parallelization

GLIMPSE uses overlapping chunks of markers to perform imputation. To define the chunks on the chromosome of interest it uses information from the reference and target panel together, keeping track of the number of variants present in each dataset. The algorithm starts by considering the whole chromosome as full chunk, then recursively divides the current chunk in half, until at least one of the following conditions is not satisfied: the imputation region (i) has at least *M*_*i*_ markers and (ii) is at least *W*_*i*_ Mb long; the buffer region (iii) has at least *M*_*b*_ markers and (iv) is at least *W*_*b*_ Mb long. If a chunk does not respect one of these conditions, then the recursion stops, and the previous chunk is returned.

We run GLIMPSE exploiting two complementary parallelization strategies to take advantage of modern computational clusters. First, we carried out imputation into overlapping chunks of data spanning on 2 Mb regions with 200kb buffer, allowing us to leverage multiple compute nodes. Second, in each chunk of data multiple samples can be processed in parallel using multi-threading, therefore taking advantage of multiple CPUs being available per node.

#### Output information

The output of GLIMPSE takes the form of a VCF/BCF file containing for each target sample the best-guess phased genotype (FORMAT/GT field), the imputed genotype posteriors averaged over all the main iterations (FORMAT/GP) and the imputed genotype dosages of the genotype posteriors (FORMAT/DS). In addition to this, GLIMPSE outputs an additional field that contains up to 16 likely haplotype pairs that have been sampled throughout the main iterations (FORMAT/HS). The HS field is a new feature of phasing methods, since it allows accurate consensus phased calls in subsequent analysis. We store the HS field in a single 32-bit integer per individual per variant.

#### Merging phased regions

Merging different imputed chunks is straightforward in the case of standard imputation methods for GWAS, where phased information is discarded. However, in the case of low-coverage imputation, is it desirable to maintain and merge phased information across several chunks. For this reason, merging together different chunks requires merging information stored in the HS field coherently. We solve this problem by minimizing Hamming distance within the overlaps between the successive phased chunks. The result is a set of phased calls across entire chromosomes.

### Benchmarks

#### Imputation performance

We measured imputation performance as the squared Pearson correlation between the validation (i.e. high-coverage) and the imputed genotypes. Since, imputation performance heavily depends on allele frequency, we measured it within multiple frequency bins that we defined using independent frequency estimates made from 71,702 fully sequenced genomes from the Genome Aggregation Database (GnomAD) version 3 ^18^. This database contains 32,299 and 21,042 genomes with Non-Finnish European and African/African-American ancestries, respectively, allowing us to have highly accurate frequency estimates even at rare variants. In practice, we used the Non-Finnish European frequencies to evaluate imputation performance of the 503 European (EUR) samples and the African/African-American frequencies for the 61 African-American (ASW) samples. From these frequency estimates, we classified all HRC variants in the overlap with GnomAD v3 into the following 17 frequency bins: ]0,0.0002], ]0.0002,0.0005], ]0.0005,0.001], ]0.001,0.002], ]0.002,0.005], ]0.005,0.01], ]0.01,0.02], ]0.02,0.05], ]0.05,0.1], ]0.1,0.15], ]0.15,0.2], ]0.2,0.25], ]0.25,0.3], ]0.3,0.35], ]0.35,0.4], ]0.4,0.45] and ]0.45,0.5]. To get reliable measures of imputation performance, we used an aggregative approach: we pooled all validation and imputed genotypes belonging to the same frequency bin together in order to compute a single squared Pearson correlation value per bin. Unless explicitly stated in the text, we measured imputation performance genome-wide across the 22 autosomes, conversely to many other benchmarks published so far that only focus on one or two chromosomes ^8-10^. As a result, we measure imputation performance at a large amount of validation data: the frequency bins contain from 25,919,013 (ASW for bin ]0.45,0.5]) to 6,326,725,930 (EUR for bin ]0,0.0002]) validation genotypes (Supplementary Table 1). Given the large amount of validation data, we developed a benchmark tool, GLIMPSE_concordance, as part of the GLIMPSE tools suite that measures the squared Pearson correlation within user-defined frequency by streaming the imputed and validation data to maintain low memory requirement (i.e. no need to store all data in memory before computing the correlations).

#### Phasing performance

We measured phasing performance as the Switch Error Rate (SER) between haplotypes estimated by GLIMPSE and Genome In A Bottle (GIAB; https://github.com/genome-in-a-bottle). GIAB defines highly accurate haplotypes for one 1,000 Genomes European individual, NA12878, that were derived from (i) available family data and (ii) long sequencing reads technologies ^19^. The SER is defined as the percentage of successive pairs of heterozygous genotypes exhibiting incorrect phase, i.e. at which alleles are not correctly co-localized on the same parental chromosome. Since imputation of low-coverage data does not necessarily lead to correct inference of heterozygous genotypes, we plotted the SER against the percentage of heterozygous genotypes being correctly called (we call discordance this percentage). The SER was measured genome-wide to compensate for the fact that only one individual was assessed. In addition, the phasing performance across multiple depths of coverage takes the form of a convex function. This is an expected result as the set of heterozygous genotypes being examined varies. With deeper coverage, more heterozygous genotypes are considered in the analysis, with lower allele frequency and shorter distance between them. As phasing is known to perform better between nearby variants and worse for rare variants, we thus obtain this convex function.

#### Low Coverage imputation methods

Over the last few years, multiple methods that allow imputation from low-coverage data have been developed. For our benchmarks, we compared GLIMPSE with three of the most commonly used imputation methods: BEAGLE v4.1 ^8^ (27Jan18.7e1), GENEIMP v1.3 ^9^, STITCH v1.6.2 ^10^. We run all imputation analysis using a window size of 2Mb, resulting in 129 chunks for chromosome 1. All the methods were run using 8 threads.

We found difficult to run low-coverage sequencing methods other than GLIMPSE, due to the unprecedented size of the reference panel used in our experiments. The only exception is for the software STITCH, which uses reference panel information only for the EM algorithm defining the set ancestral haplotypes.

##### BEAGLE v4.1

We first ran BEAGLE v4.1 with default settings and obtained prohibitive running for all the reference panels tested. Therefore, as performed we increased the modelscale parameter to a value of 2.0 and the number of phasing iterations to 0. The modelscale parameter specifies the scale of the model when sampling haplotypes for unrelated individuals (default value 0.8). The BEAGLE authors specify that increasing this parameter will trade reduced phasing accuracy for reduced runtime in the context of SNP array data, but when estimating posterior probabilities from genotype likelihoods, increasing the modelscale parameter could improve both accuracy and run-time. To run the method, we also set a maximum heap size to 32GB (Xmx32G) and increased the stack size to 16Mb (-Xss16m).

##### GENEIMP v1.3

In order to run GENEIMP on the same 2Mb windows as the other methods, we manually split the reference and target panel into chunks of 2Mb (plus a buffer region at the borders). For 1x coverage data, we used the following parameters: klthresh=40, flanksize=0.1, numfilterhaps=200, filtermethod=pairrand. The klthresh parameter is dependent on the coverage used, the flanksize parameter was set to 0.1 to have approximately an agreement in terms of buffer size to other methods and the other represent default parameters. The first step of GENEIMP is to transform the reference panel into a binary representation stored on disk, by using the bigmemory R package. This allows GENEIMP to use RAM, when there is enough RAM available, but can also create file-backed data structures, which are accessed in a fast manner when not enough RAM is available. However, we found that the creation of this binary data structure, which is a first mandatory step of the algorithm, to be highly RAM expensive and big reference panels can hardly be stored in this representation. For example, storing the full HRC chromosome 1 in this file format, would need more than 20 TB of disk space. In addition to this, while we found that by using file-backed data structures the imputation step is light in memory, the creation of the reference panel data structure requires hundreds of GB for a single chunk. For this reason, we were only able to run GENEIMP up to 5000 samples in the reference panel. We tried different values of the klthresh parameter, but we were only able to complete chromosome-wide imputation for 1.0x coverage using klthresh=40 and all other coverages ended with errors messages. When all the imputation jobs for the chromosomes ended, we removed buffer regions from each of the imputed chunks and merged the chunks together using bcftools.

##### STITCH v1.6.2

We run STITCH with and without reference panel information. In the STITCH method the reference panel is only used for initialisation in the Expectation Maximization algorithm used to estimate the parameters of the hidden Markov model, and not for subsequent updating iterations.

The most important parameter in the STITCH model is the number of founder haplotypes the model uses, indicated as *K*. In our tests we used a value of *K* = 10 since it is reasonable to keep the computational cost of the diploid model low. We also used *K* = 40 for few configurations to verify the effect of the *K* parameter (Supplementary Figure 18). For both the values of *K*, we set the parameter *nGen* = 4*N*_*e*_/*K*, where *N*_*e*_ has been set to have a value of 20,000 as recommended by the STITCH authors. STITCH uses the BAM files directly and not the genotype likelihoods, differently from other methods. We found that the method depends heavily on the choice of *K*, but there is no clear best strategy of the value to use, since the gain at rare variants obtained using *K* = 40 (with a computational overhead) is only visible for higher coverage settings (≥ 1x) and is compensated by a loss in accuracy at common variants, which is mainly the focus of the method.

We point out that STITCH is designed for imputation without a reference panel, especially for non-human species. For human samples STITCH can be used to capture variation at common variants. However, its performance drops considerably compared to reference-based approaches at rare variants, even if the reference panel does not represent the target population particularly well, in terms of genetic ancestry.

#### SNP array imputation methods

Over the last decade, multiple methods able to impute SNP array data have been developed ^8,12,15,20^. For our benchmarks, we compared GLIMPSE with two different generations of methods: BEAGLE v4.1 (27Jan18.7e1) and BEAGLE v5.1 (25Nov19.28d). Both methods perform pre-phasing and imputation. We run imputation genome-wide (chromosome 1 to 22) for the 25 simulated SNP array datasets using default imputation window size as defined in BEAGLE v4.1 and v5.1. Both methods were run using 8 threads. For BEAGLE v4.1, in order to run the pre-phasing step using HRC as a reference panel, we had to change some of the default parameter values. Specifically, we used the same parameter setting than in the case of imputation from low-coverage sequencing data: *modelscale*=2.0 and *niterations*=0. Concerning the imputation step, we used default settings for all parameters and ran imputation for each of the 22 autosomes separately. In contrast, BEAGLE v5.1 was run using default values for all parameters.

### Mapping expression quantitative trait loci

We identified expression QTLs (eQTLs) for each quantified gene using the QTLtools v1.1 software ^27^. Briefly, this comprises performing (i) 1000 permutations to correct for the number of genetic variants being tested per gene in cis (±1 Mb from the gene Transcription Start sites) and (ii) apply a false discovery rate correction (FDR 5%) using the Benjamini-Hochberg procedure to correct for the number of genes being tested genome-wide. For both low-coverage and SNP array datasets, only genetic variants with a minor allele frequency (directly computed from the genotype dosages) above 1% were used. We identified eGenes (genes at which expression is under genetic control) in each dataset by selecting those being associated with at least one genetic variant (FDR 5%). The accuracy, precision and recall of eGene discovery per dataset was calculated using eGenes identified with high-coverage as the ground truth. For the comparison of p-values between high-coverage and each dataset for the same genetic variants the set of lead eQTLs obtained for the high-coverage dataset were used. P-values were −log10-transformed and the value discrepancy between the ground truth (high-coverage) and each dataset was measured through mean absolute error. The lead eQTL of each eGene is defined as the cis variant with the lowest p-value of association. When more than one variant is associated to the same gene with the same exact p-value (e.g. due to complete linkage disequilibrium), one of these variants is randomly chosen.

The overlap between the coordinates of lead eQTLs identified in each dataset with functional regions was obtained using the pybedtools python library ^30^. Coordinates of LCL-specific (i) protein binding sites (from multiple DNA binding proteins), (ii) DNase I–hypersensitive sites (DNAse-seq; narrow peaks) and (iii) locations with H3K27ac histone modifications (ChIP-seq; narrow peaks from two replicates) were downloaded from the ENCODE project (integration_data_jan2011, ENCSR000EJD and ENCSR000AKC, respectively). Protein binding sites were lifted from hg19 to hg38 using the UCSC liftover tool.

### Burden test analysis

For the burden test analysis in coding regions, we used gene and exon coordinates given by the Gencode annotation v33 ^26^ for autosomal chromosomes. Specifically, we extracted all annotated exons (n=1,108,031 exons) belonging to all protein coding genes (n=19,080 genes). Then, to avoid double counting some rare variants, we merged all overlapping exons into non-overlapping exonic regions (n=225,581). Finally, since some of these exonic regions belong to multiple genes, we ended up with a total of 234,411 exonic regions assigned to 19,080 genes; meaning that we have on average 12.3 (sd=11.5) exonic regions per gene. The burden of rare variants was computed as follows for each call set: (i) we extracted all variants with a MAF below 1% that fall within the exonic regions, (ii) we summed the minor allele dosages at these variants per individual and per gene. Of note here, both the MAF and the minor allele dosages are computed from the allelic dosages obtained post-imputation (VCF/DS field). Finally, this procedure has been done for the high-coverage call set and all other call sets either imputed from low-coverage sequencing or SNP arrays. Then, as a measure of imputation accuracy, we measured the squared Pearson correlation between the burdens derived from high-coverage and those from imputation.

## Data availability

GLIMPSE tools: https://github.com/odelaneau/GLIMPSE Website: https://odelaneau.github.io/GLIMPSE/

## Acknowledgments

The NYGC 1000 Genomes data were generated at the New York Genome Center with funds provided by NHGRI Grant 3UM1HG008901-03S1. This work was funded by a Swiss National Science Foundation (SNSF) project grant (PP00P3_176977)

## Author contribution

S.R., D.M.R. and O.D. designed the study, performed experiments and drafted the paper. S.R. and O.D. developed the algorithm and wrote the software. S.R., R.H. and O.D. created the website. O.D. supervised the project. All authors reviewed the final manuscript.

## Corresponding author

Olivier Delaneau

