## Supplementary Information for "Efficient phasing and imputation of low-coverage sequencing data using large reference panels"

10/04/2020

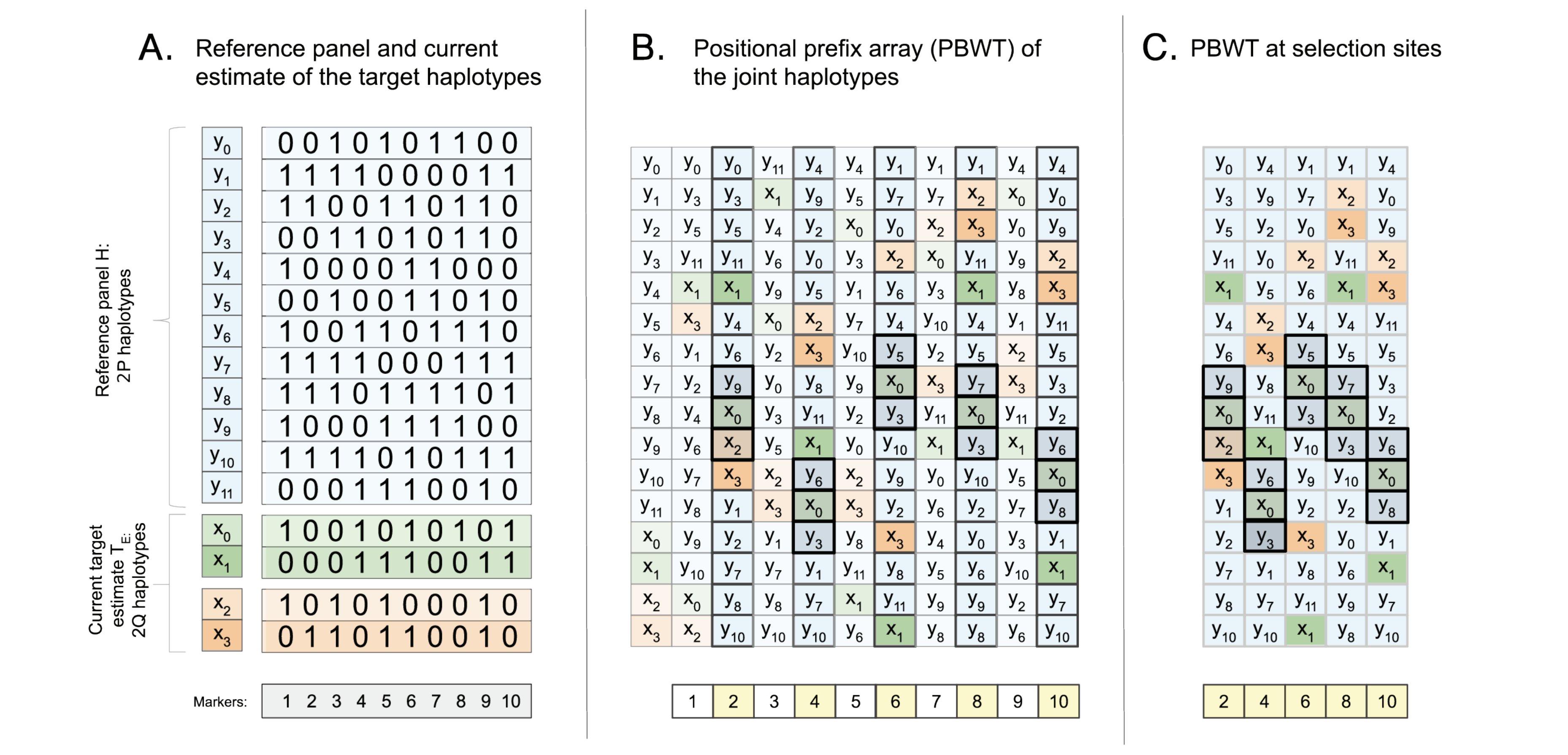


**<u>Supplementary Figure 1:</u> PBWT state selection.** (**A**) The GLIMPSE state selection takes information from a reference panel of haplotypes (light blue) and the current haplotype estimate of the target individuals (green for individual 1 and orange for individual 2). (**B**) The joint positional prefix array is created at each marker. Haplotypes that share a reverse prefix cluster together in the data structure. Here, only 1 neighbour in each direction is shown (pbwt-depth=1) for haplotype $x_0$, every 2 markers (pbwt-modulo=2) (dark grey colour). (**C**) Zoom-in view at selection sites. At the end of the selection step, the list of selected haplotypes is: $\{x_2, x_3, x_5, x_6, x_7, x_8, x_9\}$. Selected states of both the haplotypes of an individual are merged together and used for phasing and imputation.

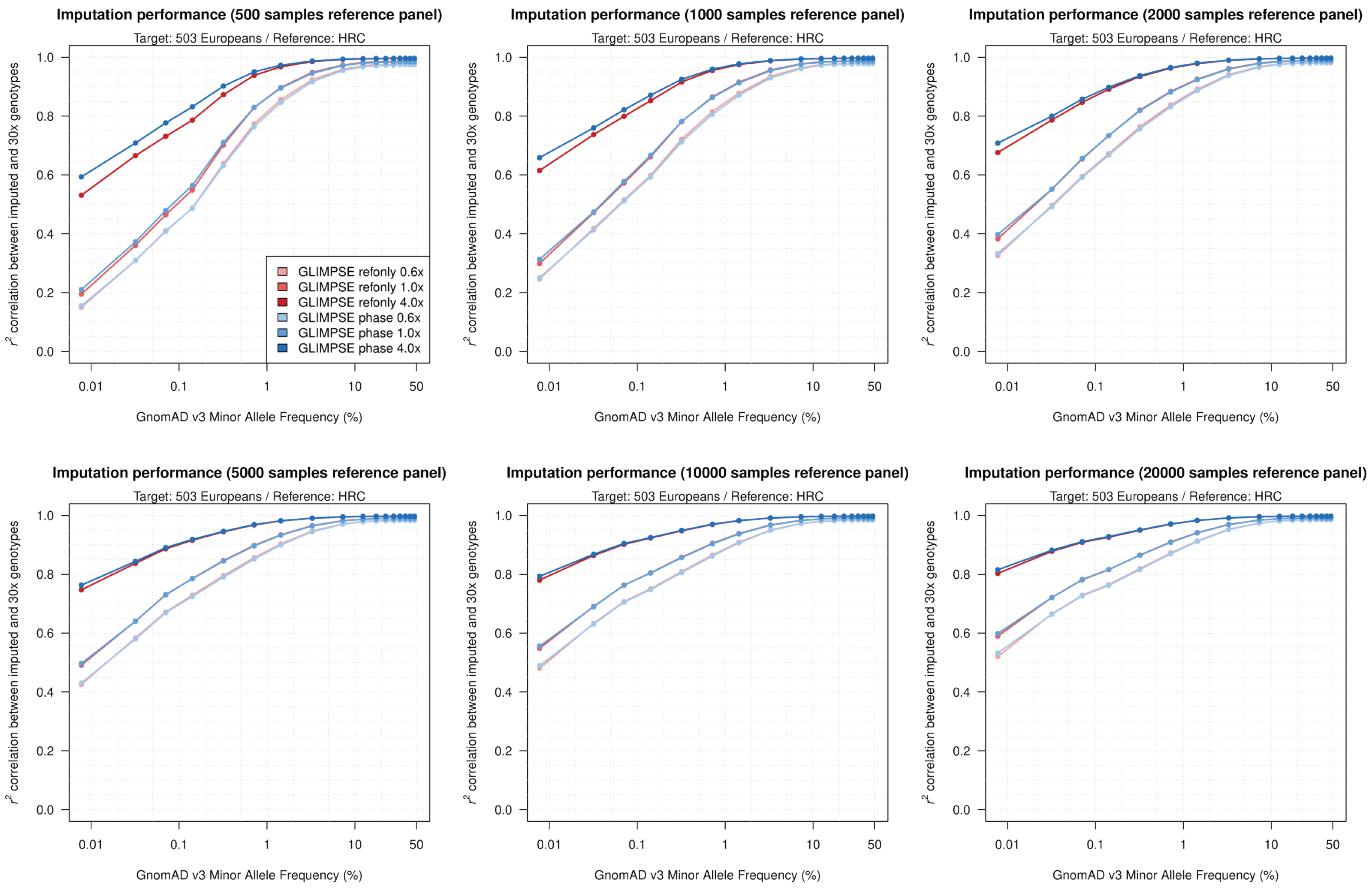


**<u>Supplementary Figure 2:</u> Imputation performance using the full conditioning set and using only reference panel haplotypes.** Imputation performance using chromosome 1 data from the coverage European population dataset (0.6x, 1x and 4x coverage) for different reference panel sizes, using the full conditioning set (blue) and using only reference panel haplotypes (red). The horizontal axis is on a log scale.

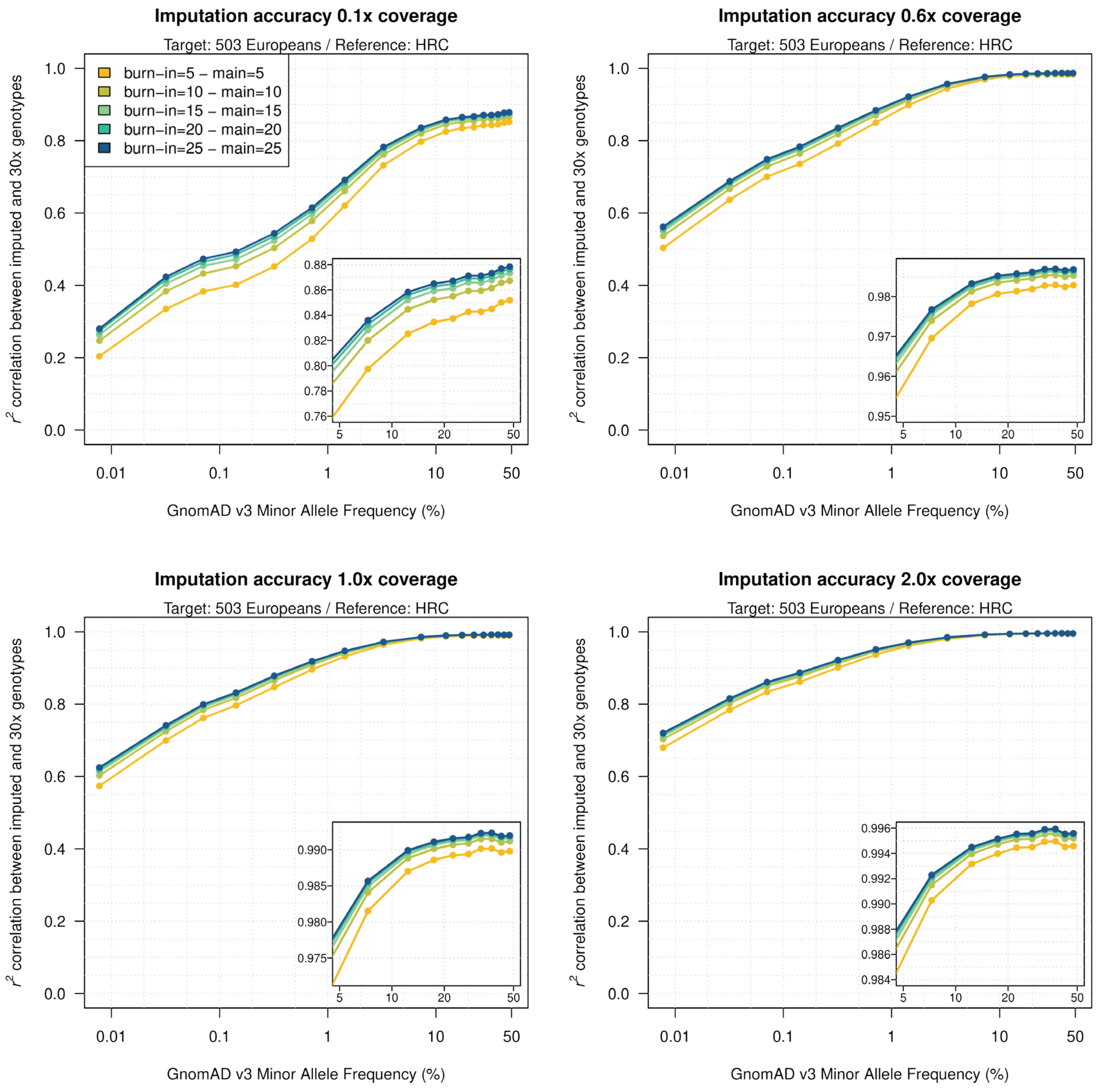


**<u>Supplementary Figure 3:</u> Imputation performance varying the number of iterations.** Imputation performance using chromosome 1 data from the European population dataset and the full HRC as a reference panel for different coverage configurations, varying the number of iterations. The horizontal axis is on a log scale. Increasing the number of iterations has a noticeable effect only for extremely low-coverage target datasets (0.1x coverage), where more iterations may be needed to reach convergence.

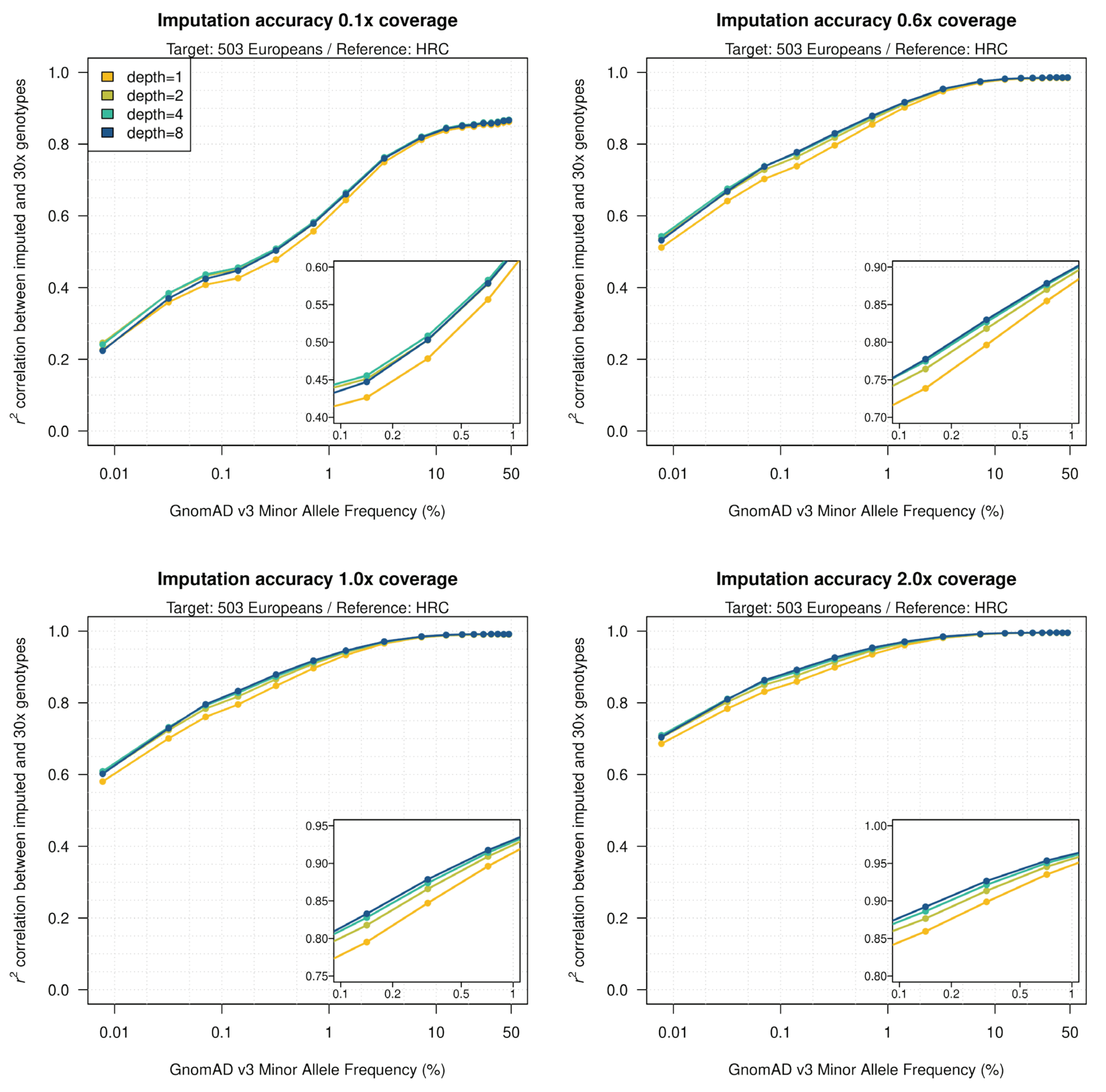


**Supplementary Figure 4: Imputation performance varying the PBWT parameters.** Imputation performance using chromosome 1 data from the European population dataset and the full HRC as a reference panel for different coverage configurations, varying the pbwt-depth parameter. The method is very robust to even extremely low values of the parameter. We notice that moderate values of the pbwt-depth seem to perform better at extremely rare variants, compared to high values. A possible explanation is that the for high value of the pbwt-depth the cap in the number of states (set to K, the number of initialising states) is reached and the sampling strategy tends to prefer long IBS states compared to locally optimal ones. The horizontal axis is on a log scale.

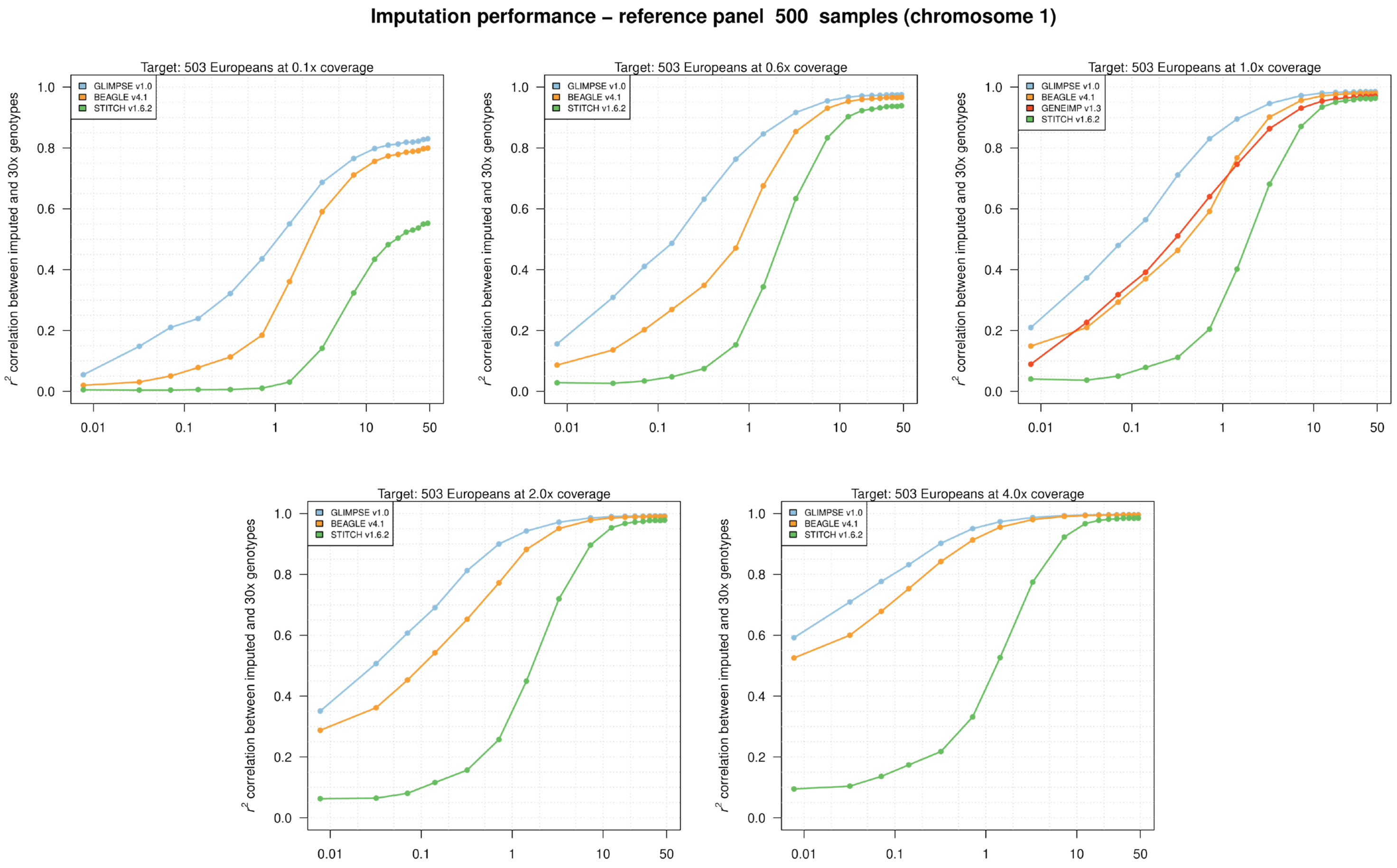


**<u>Supplementary Figure 5:</u> Imputation performance using a reference panel containing 500 samples.** Imputation performance of different low-coverage imputation methods using chromosome 1 data from the European population dataset, a subset of 500 samples from the HRC as a reference panel, varying the sequencing coverage of the target dataset. The horizontal axis is on a log scale. We only ran GENEIMP on 1x coverage data (see **Online methods**).

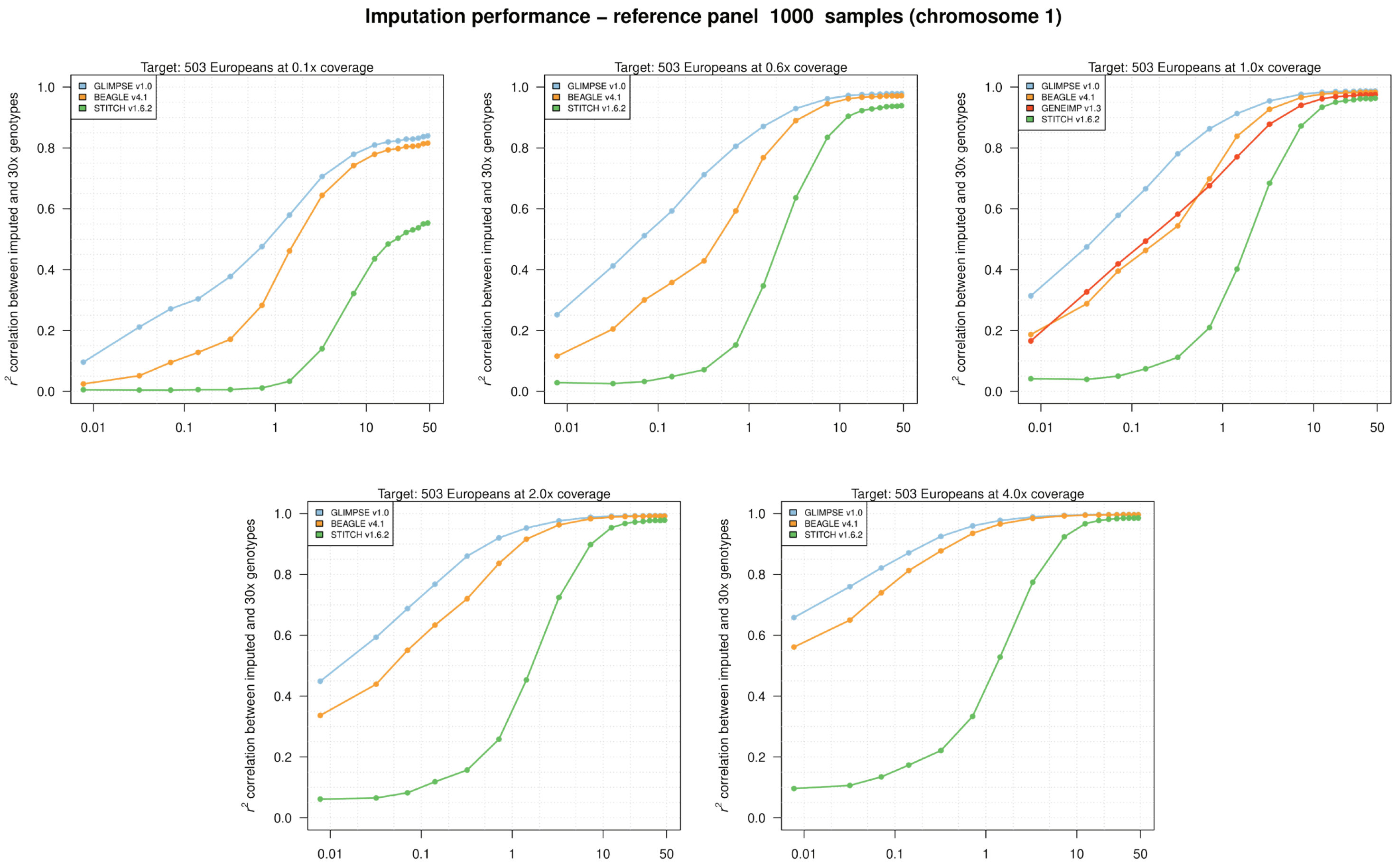


**<u>Supplementary Figure 6:</u> Imputation performance using a reference panel containing 1,000 samples.** Imputation performance of different low-coverage imputation methods using chromosome 1 data from the European population dataset, a subset of 1,000 samples from the HRC as a reference panel, varying the sequencing coverage of the target dataset. The horizontal axis is on a log scale. We only ran GENEIMP on 1x coverage data (see **Online methods**).

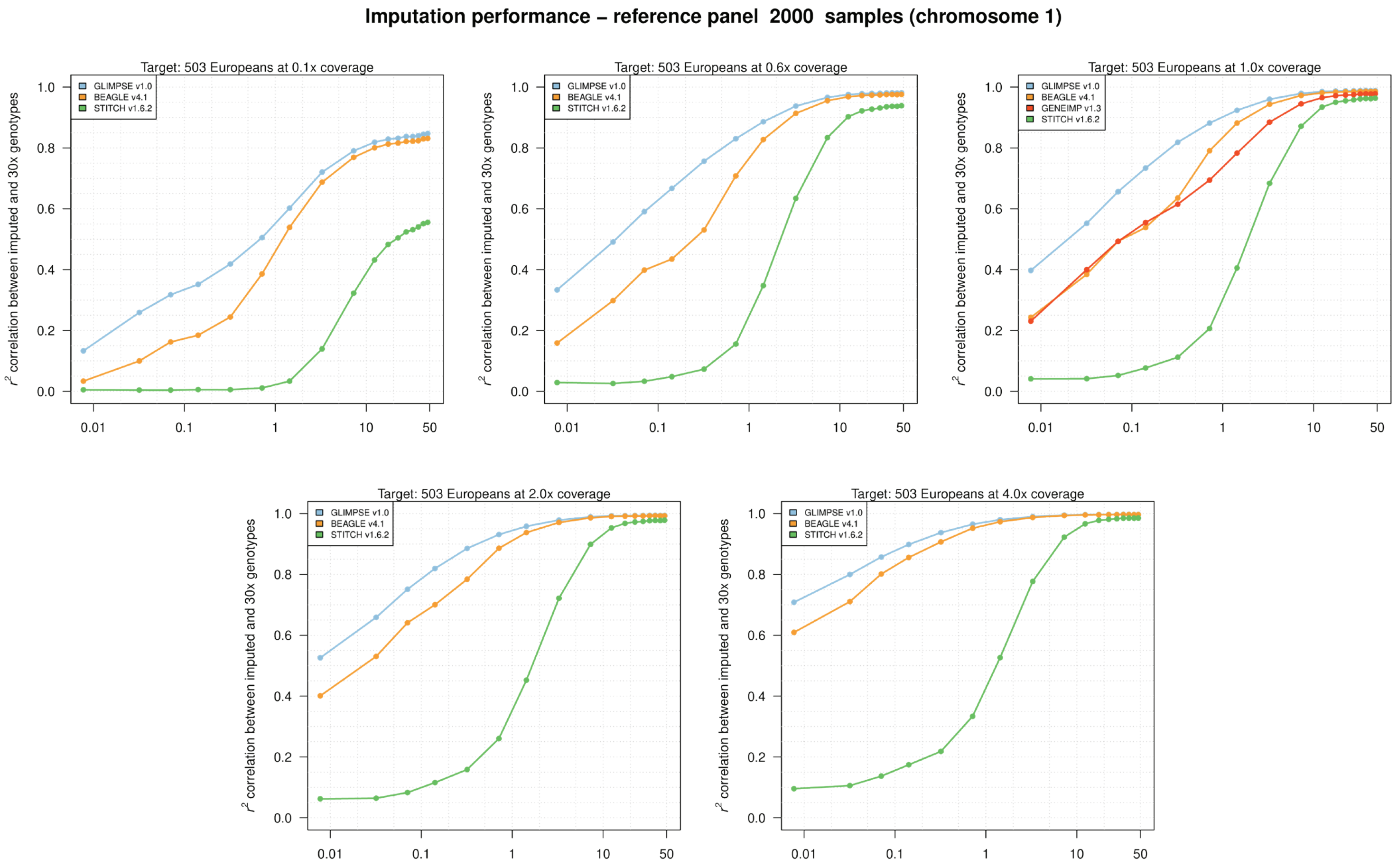


**<u>Supplementary Figure 7:</u> Imputation performance using a reference panel containing 2,000 samples.** Imputation performance of different low-coverage imputation methods using chromosome 1 data from the European population dataset, a subset of 2,000 samples from the HRC as a reference panel, varying the sequencing coverage of the target dataset. The horizontal axis is on a log scale. We only ran GENEIMP on 1x coverage data (see **Online methods**).

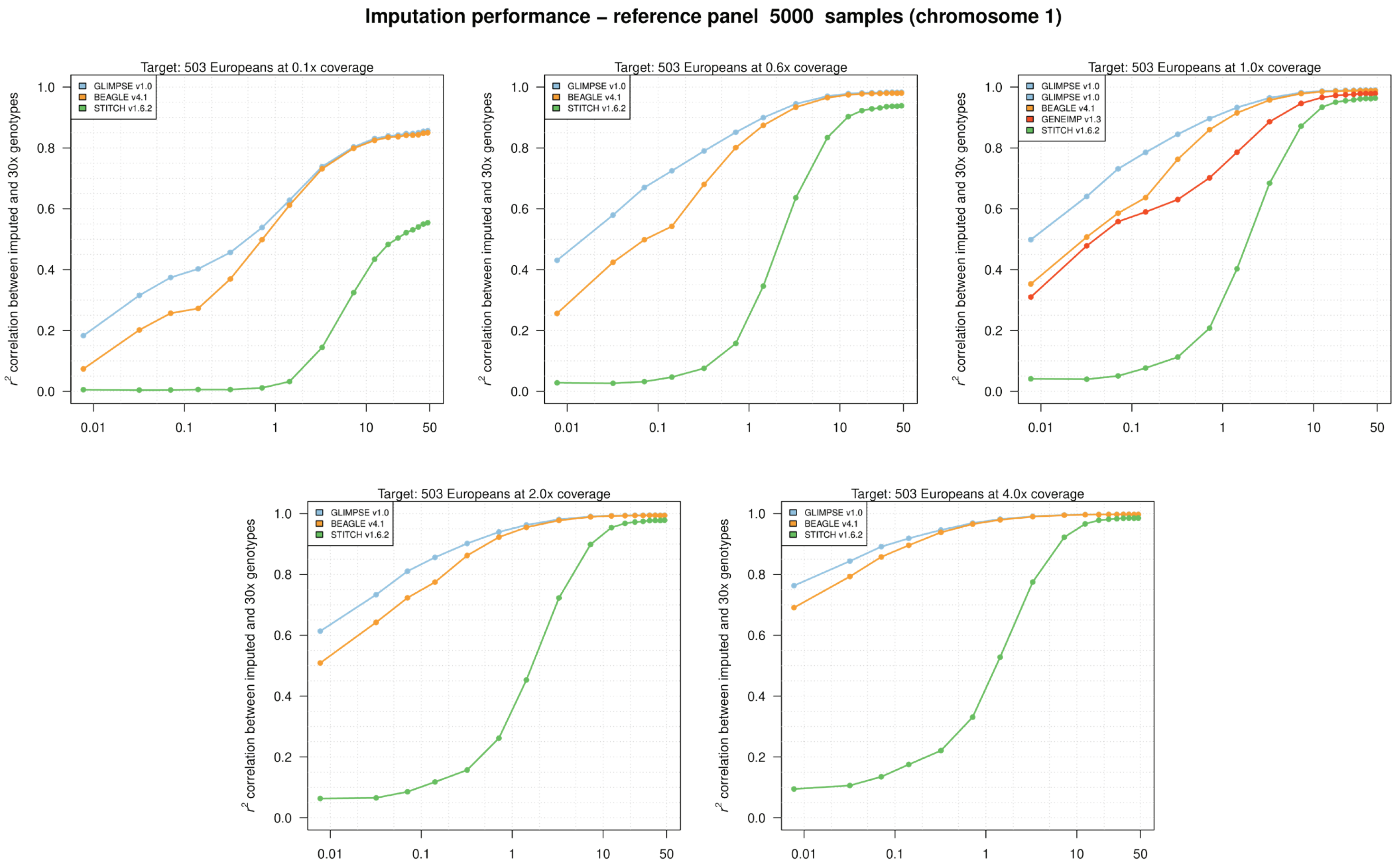


**<u>Supplementary Figure 8:</u> Imputation performance using a reference panel containing 5,000 samples.** Imputation performance of different low-coverage imputation methods using chromosome 1 data from the European population dataset, a subset of 5,000 samples from the HRC as a reference panel, varying the sequencing coverage of the target dataset. The horizontal axis is on a log scale. We only ran GENEIMP on 1x coverage data (see **Online methods**).

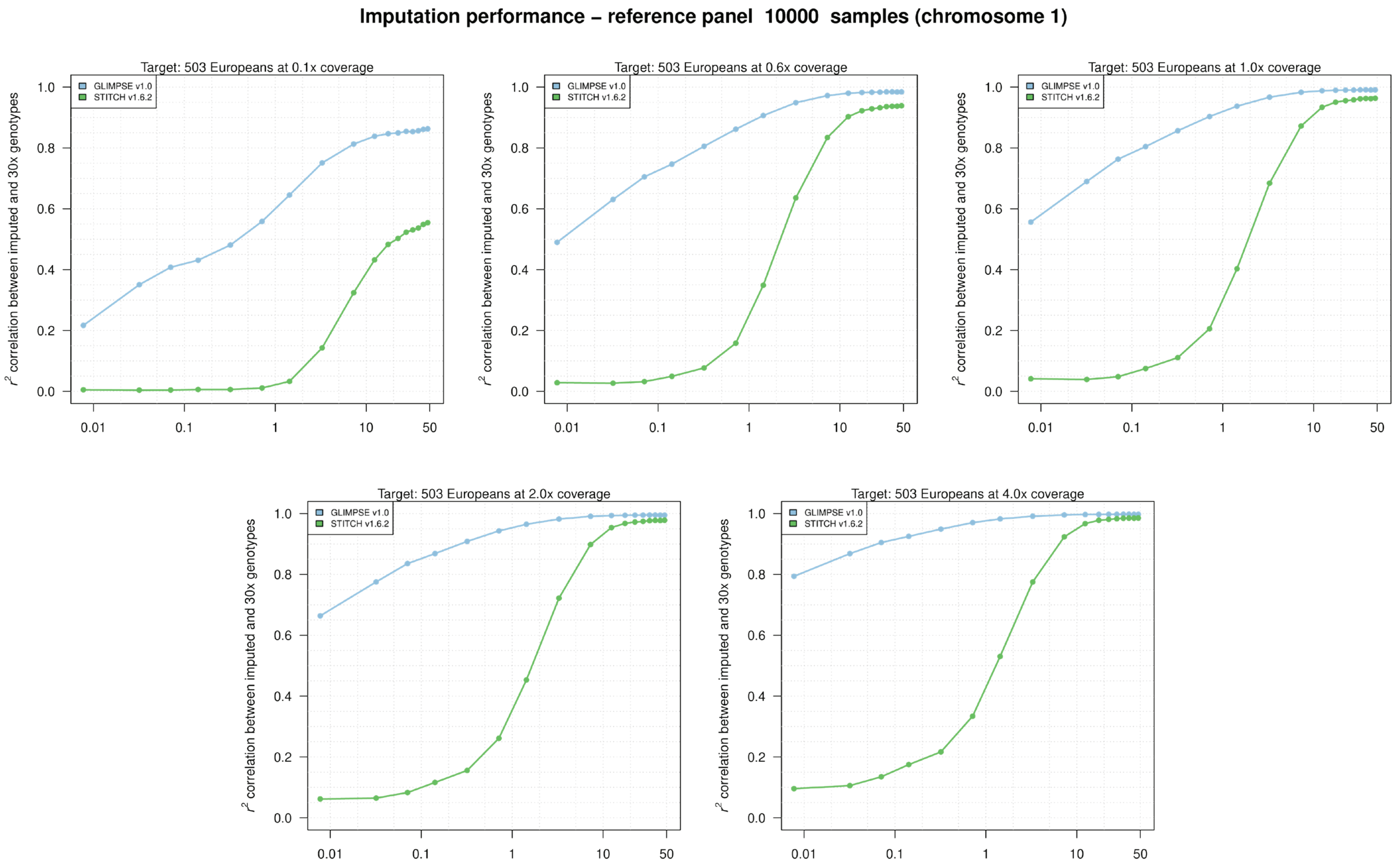


**<u>Supplementary Figure 9:</u> Imputation performance using a reference panel containing 10,000 samples.** Imputation performance of different low-coverage imputation methods using chromosome 1 data from the European population dataset, a subset of 10,000 samples from the HRC as a reference panel, varying the sequencing coverage of the target dataset. The horizontal axis is on a log scale. We were not able to run GENEIMP and BEAGLE on this reference panel, due to time limits.

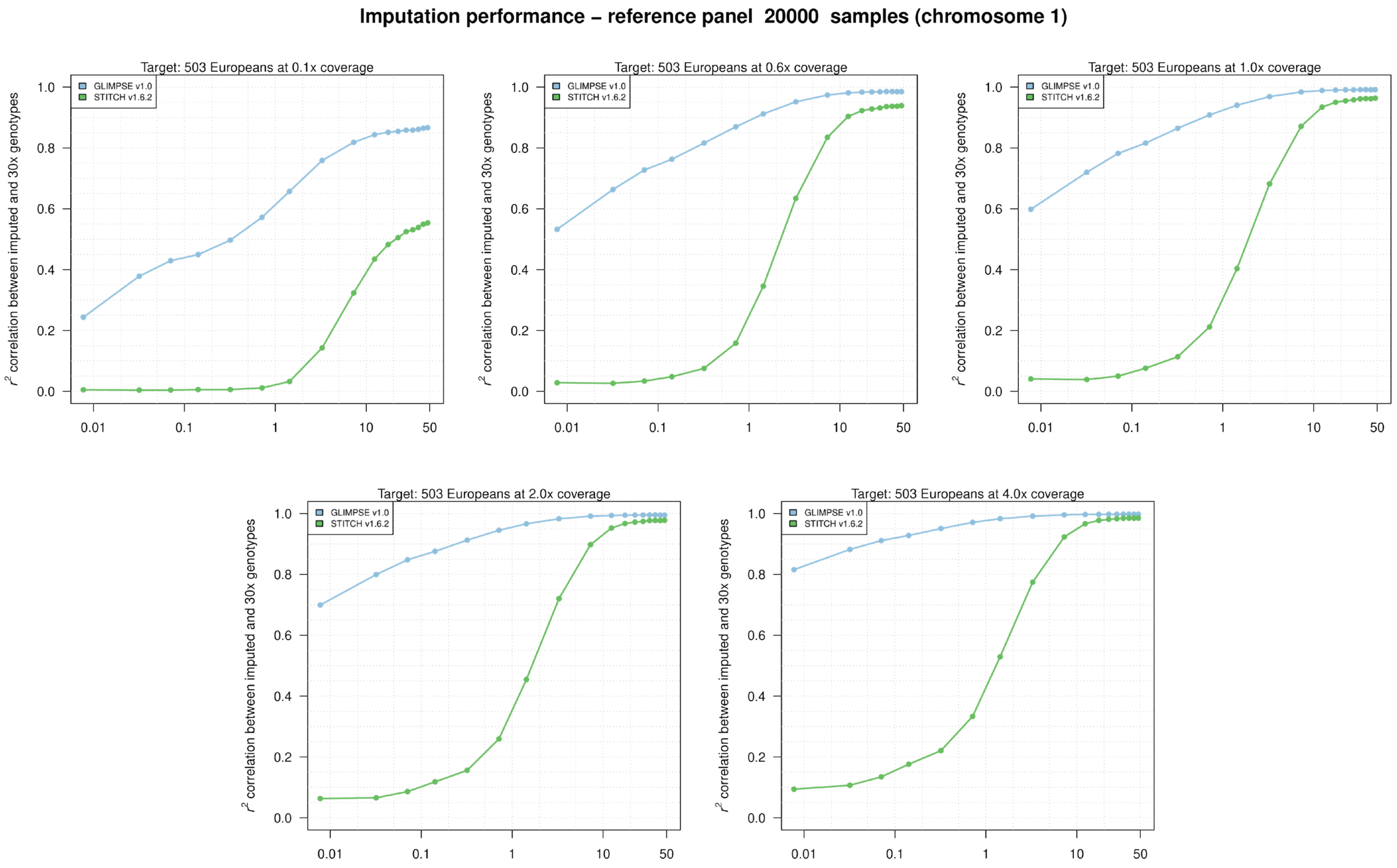


**<u>Supplementary Figure 10:</u> Imputation performance using a reference panel containing 20,000 samples.** Imputation performance of different low-coverage imputation methods using chromosome 1 data from the European population dataset, a subset of 20,000 samples from the HRC as a reference panel, varying the sequencing coverage of the target dataset. The horizontal axis is on a log scale. We were not able to run GENEIMP and BEAGLE on this reference panel, due to time limits.

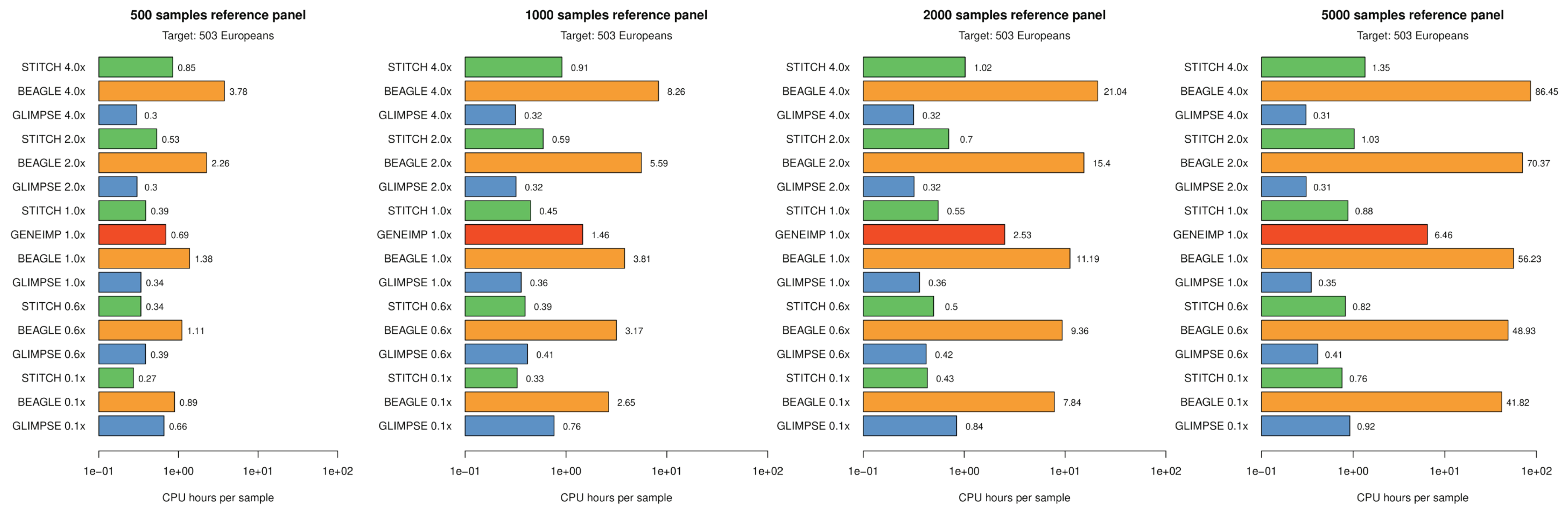


**Supplementary Figure 11. Running times of different imputation methods.** Imputation performance of low-coverage sequencing methods for the European population chromosome 1 dataset. We were only able to run GENEIMP using 1x coverage data. We only show reference panel size up to 5,000 samples due to time limits. The horizontal axis is on a log scale.

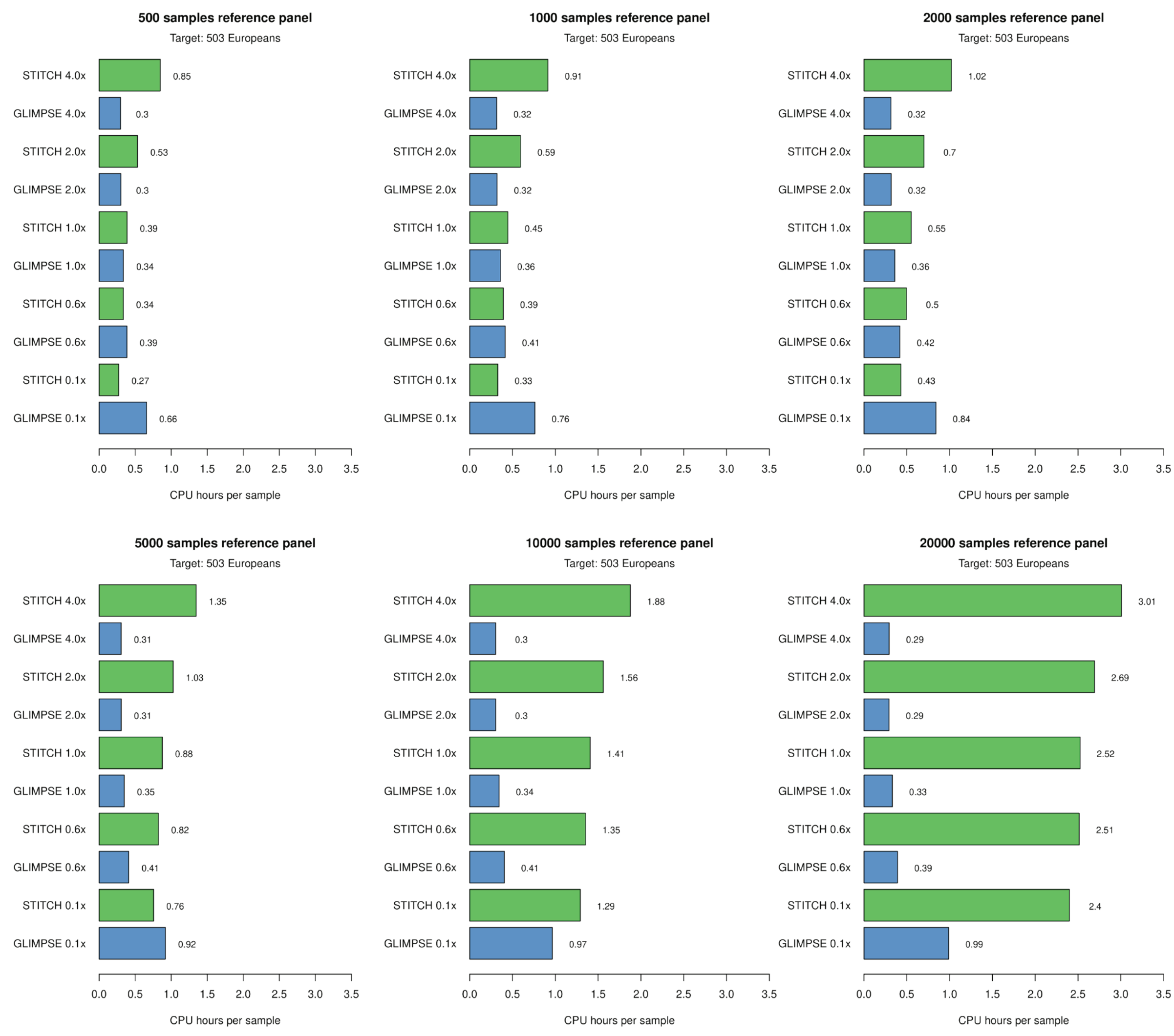


**Supplementary Figure 12: Running times of GLIMPSE and STITCH for the full set of downsampled reference panels.** Imputation performance of GLIMPSE and STITCH for the European population chromosome 1 dataset on the full set of downsampled reference panels.

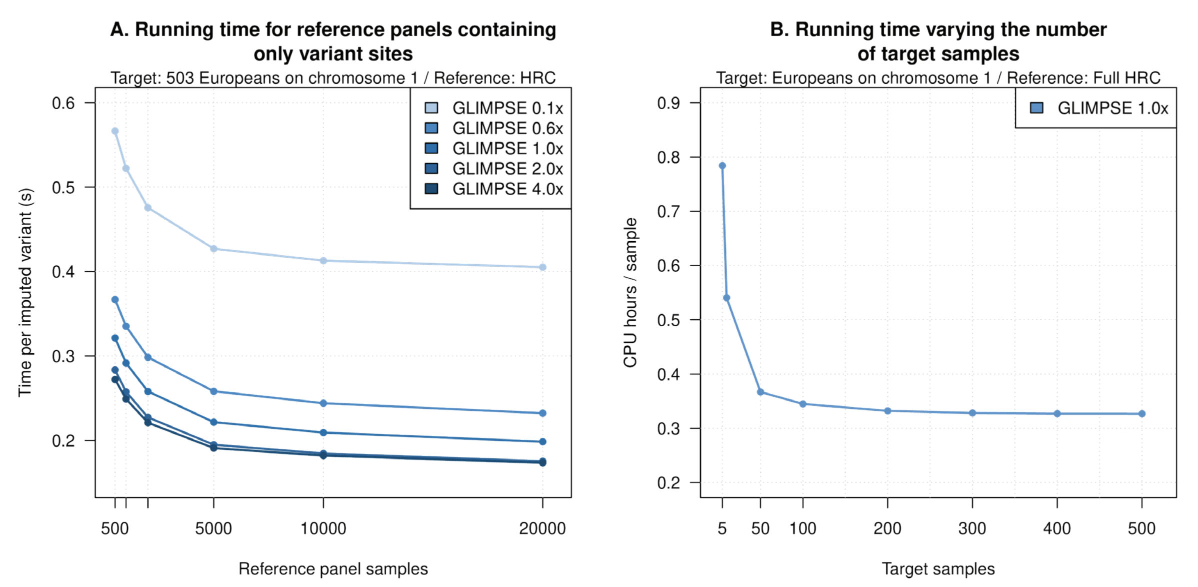


**Supplementary Figure 13: Running times versus target samples.** (**A**) Running time of GLIMPSE to impute a single variants (y-axis) on chromosome 1 using multiple coverages of the European population dataset (shades of blues) and multiple subsets of the reference panel (x-axis), only retaining variable positions. (**B**) Running times of GLIMPSE (y-axis) to impute the whole chromosome 1 using HRC as reference and an increasing number of target samples (x-axis).

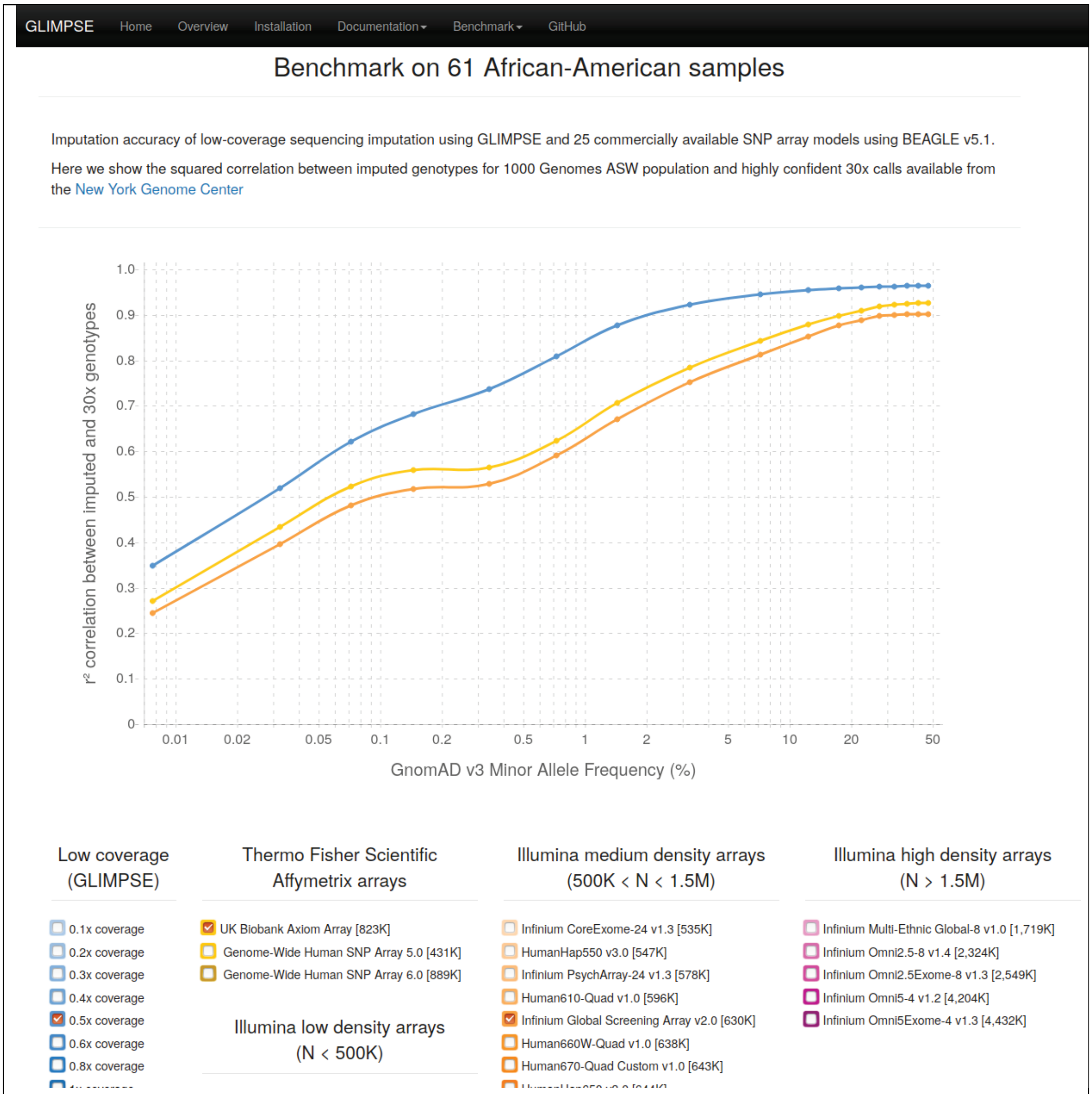


**Supplementary Figure 14: GLIMPSE website screenshot.** https://odelaneau.github.io/GLIMPSE/

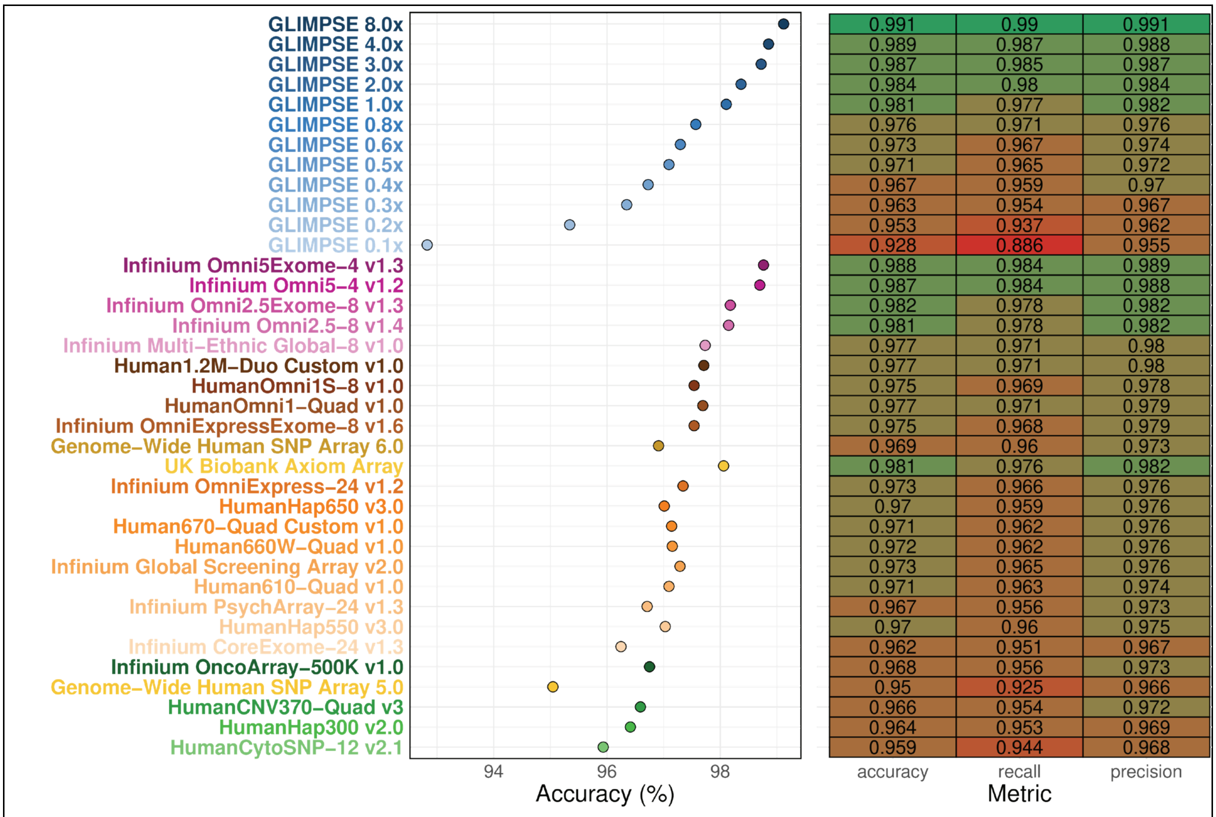


**<u>Supplementary Figure 15:</u> eGene identification accuracy, recall and precision.** Accuracy, recall and precision of eGene identification between each dataset and high-coverage (30x), the latter used as the ground truth for these metrics.

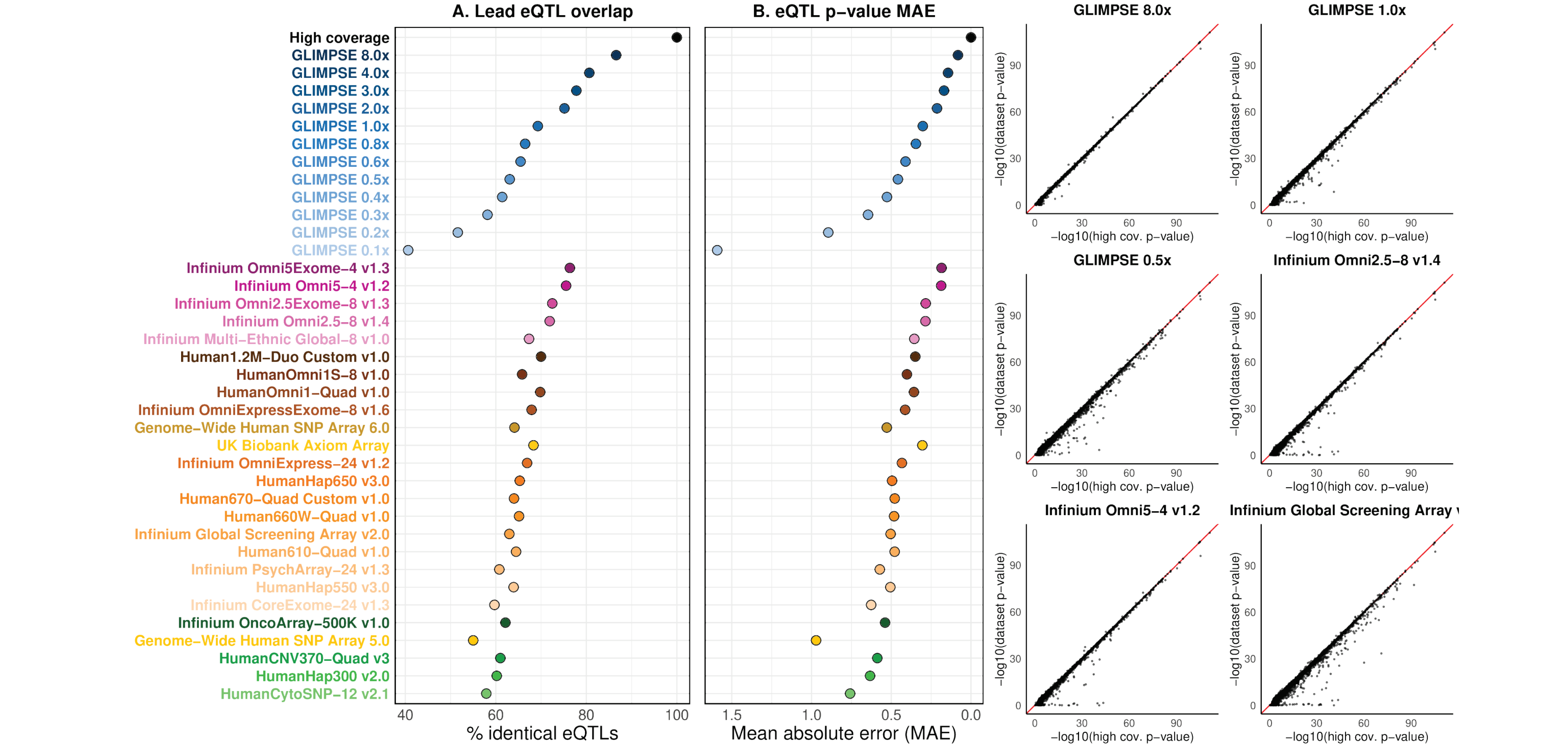


**Supplementary Figure 16: Lead eQTL overlap and association p-value mean absolute error. (A)** Overlap between lead eQTLs identified in high-coverage and each low-coverage and SNP array dataset. eQTL mapping was performed independently for each dataset (FDR 5%; MAF >= 1%). eGenes in which the lead eQTL p-value was tied with another variant's p-value (e.g. due to perfect linkage disequilibrium) were excluded, as the choice of variant for being the lead eQTL in these cases is arbitrary. The total number genes assessed after filtering was 5016. **(B)** Mean absolute error between -$\log_{10}$ p-values of association obtained for high-coverage lead eQTLs and those obtained in each dataset for the same set of variants. All high-coverage lead eQTLs (i.e. a variant for each of the 16894 genes) were considered here, regardless of significance level. The scatterplots detail the -$\log_{10}$ p-values used to calculate the mean absolute errors for several relevant low-coverages and SNP arrays.

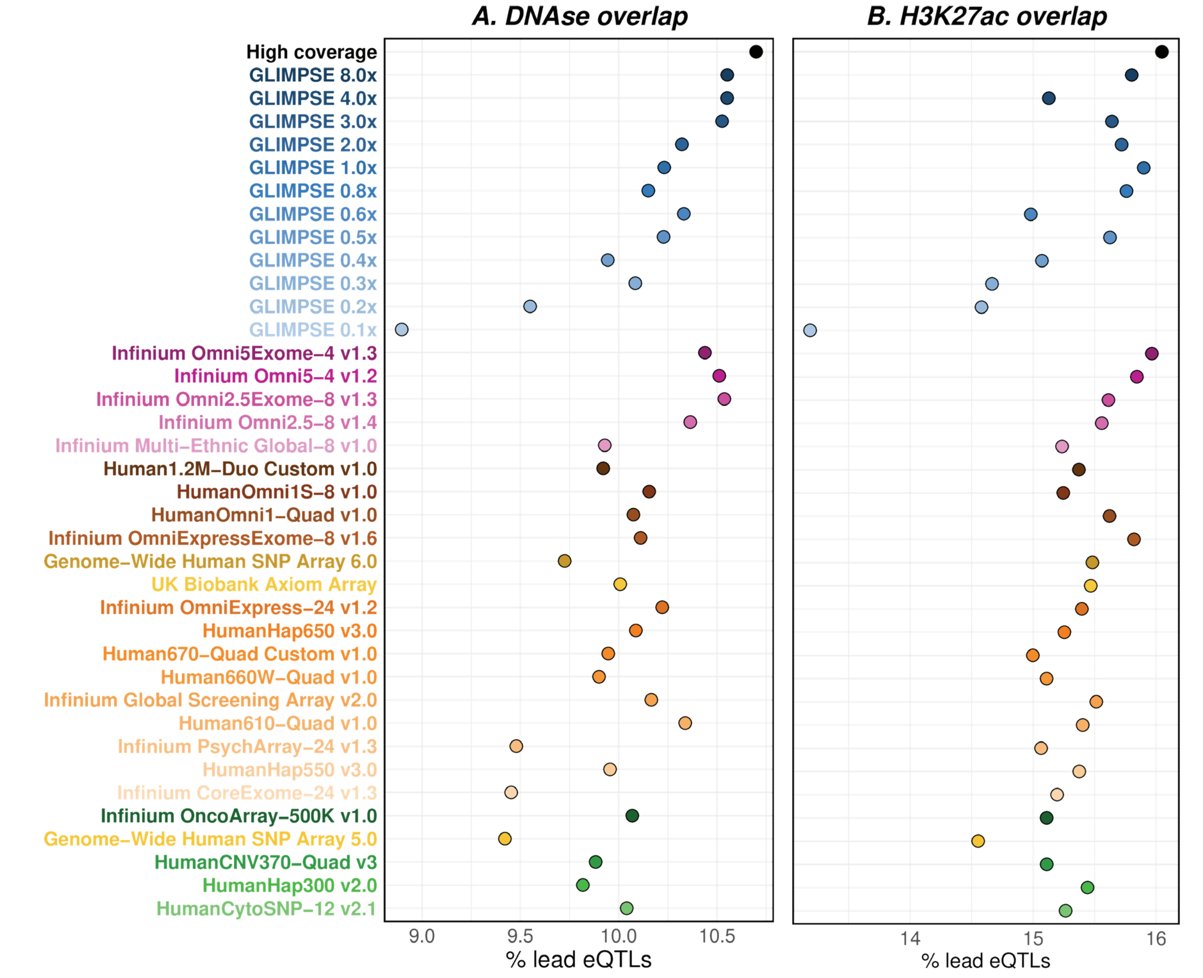


**__Supplementary Figure 17:__ Lead eQTL overlap to functional regions. (A)** Percentage of lead eQTLs overlapping DNAse regions (suggestive of open chromatin). **(B)** Percentage of lead eQTLs overlapping H3K27ac histone epigenetic modification sites (suggestive of enhancer activity). Both functional annotations come from experiments performed in LCL (GM12878). Only lead eQTLs for eGenes (i.e. significantly associated gene-variant pairs at FDR 5%) were used here.

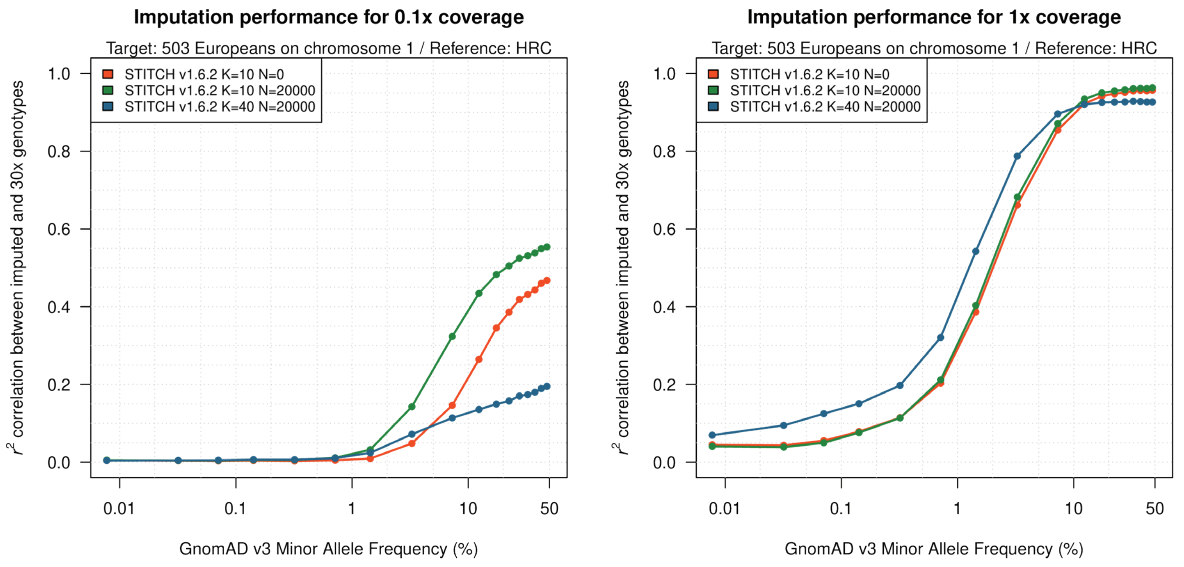


**Supplementary Figure 18: STITCH imputation performance.** Imputation performance of the STITCH method on chromosome 1 data from the European population dataset (0.1x and 1x coverage). STITCH with and without a reference panel (N=20,000 and N=0, respectively) and using the number of ancestral haplotypes (K) equal to 10 and 40. The reference panel used is the downsampled HRC containing 20,000 samples. The horizontal axis is on a log scale.

| | European population | | African-American population | |
|---|---|---|---|---|
| Index of frequency bin | Total number of validation genotypes | Mean minor allele frequency | Total number of validation genotypes | Mean minor allele frequency |
| 1 | 6,326,725,930 | 0.000078 | 519,752,417 | 0.000077 |
| 2 | 3,052,041,520 | 0.000318 | 157,037,210 | 0.000324 |
| 3 | 1,484,071,007 | 0.000706 | 96,076,590 | 0.000719 |
| 4 | 1,002,944,675 | 0.001414 | 90,390,681 | 0.001452 |
| 5 | 938,463,551 | 0.003202 | 161,625,388 | 0.003406 |
| 6 | 570,382,536 | 0.007141 | 168,647,279 | 0.007256 |
| 7 | 514,665,967 | 0.014395 | 167,205,341 | 0.014375 |
| 8 | 661,157,544 | 0.032845 | 205,462,164 | 0.032512 |
| 9 | 561,195,175 | 0.072814 | 141,023,616 | 0.071869 |
| 10 | 392,074,043 | 0.123978 | 77,367,994 | 0.123226 |
| 11 | 319,355,609 | 0.174361 | 54,630,485 | 0.173785 |
| 12 | 280,697,856 | 0.224477 | 42,628,232 | 0.224107 |
| 13 | 252,921,365 | 0.274683 | 35,450,082 | 0.274372 |
| 14 | 238,661,910 | 0.324813 | 31,198,189 | 0.324572 |
| 15 | 228,719,945 | 0.374863 | 28,373,590 | 0.374604 |
| 16 | 218,184,693 | 0.424965 | 26,220,153 | 0.424869 |
| 17 | 217,020,054 | 0.474900 | 25,919,013 | 0.474981 |

**Supplementary Table 1: Amount of validation genotypes to measure imputation performance.**

| Company | Name | #SNPs | #SNPs in HRC | %SNPs in HRC | %missing |
|---|---|---|---|---|---|
| Affymetrix | UK Biobank Axiom Array | 822,749 | 736,190 | 89.48 | 0.019 |
| Affymetrix | Genome-Wide Human SNP Array 5.0 | 430,865 | 423,351 | 98.26 | 0.012 |
| Affymetrix | Genome-Wide Human SNP Array 6.0 | 888,572 | 872,123 | 98.15 | 0.014 |
| Illumina | Infinium Global Screening Array v2.0 | 630,234 | 566,809 | 89.94 | 0.023 |
| Illumina | Human1.2M-Duo Custom v1.0 | 1,190,571 | 1,050,335 | 88.22 | 0.029 |
| Illumina | Human610-Quad v1.0 | 596,493 | 565,964 | 94.88 | 0.02 |
| Illumina | Human660W-Quad v1.0 | 638,350 | 564,854 | 88.49 | 0.038 |
| Illumina | Human670-Quad Custom v1.0 | 643,398 | 567,939 | 88.27 | 0.038 |
| Illumina | HumanCNV370-Quad v3 | 357,855 | 330,306 | 92.3 | 0.022 |
| Illumina | HumanCytoSNP-12 v2.1 | 276,462 | 263,626 | 95.36 | 0.016 |
| Illumina | HumanHap300 v2.0 | 309,143 | 305,447 | 98.8 | 0.015 |
| Illumina | HumanHap550 v3.0 | 547,472 | 540,967 | 98.81 | 0.016 |
| Illumina | HumanHap650 v3.0 | 644,265 | 636,591 | 98.81 | 0.016 |
| Illumina | HumanOmni1-Quad v1.0 | 1,072,290 | 955,623 | 89.12 | 0.027 |
| Illumina | HumanOmni1S-8 v1.0 | 1,159,020 | 1,027,562 | 88.66 | 0.037 |
| Illumina | Infinium CoreExome-24 v1.3 | 535,257 | 407,658 | 76.16 | 0.014 |
| Illumina | Infinium Omni2.5-8 v1.4 | 2,323,515 | 2,126,622 | 91.53 | 0.026 |
| Illumina | Infinium Omni2.5Exome-8 v1.3 | 2,548,888 | 2,234,605 | 87.67 | 0.025 |
| Illumina | Infinium Omni5-4 v1.2 | 4,204,349 | 3,832,684 | 91.16 | 0.023 |
| Illumina | Infinium Omni5Exome-4 v1.3 | 4,431,723 | 3,919,578 | 88.44 | 0.023 |
| Illumina | Infinium OmniExpress-24 v1.2 | 693,967 | 678,929 | 97.83 | 0.014 |
| Illumina | Infinium OmniExpressExome-8 v1.6 | 936,916 | 819,565 | 87.47 | 0.014 |
| Illumina | Infinium PsychArray-24 v1.3 | 578,434 | 435,337 | 75.26 | 0.015 |
| Illumina | Infinium Multi-Ethnic Global-8 v1.0 | 1,718,552 | 1,314,034 | 76.46 | 0.034 |
| Illumina | Infinium OncoArray-500K v1.0 | 484,284 | 454,487 | 93.85 | 0.017 |

**Supplementary Table 2:** **List of SNP arrays used in the benchmark.**
